# Frequency dependent responses of neuronal models to oscillatory inputs in current versus voltage clamp

**DOI:** 10.1101/515510

**Authors:** Horacio G. Rotstein, Farzan Nadim

**Affiliations:** Federated Department of Biological Sciences, New Jersey Institute of Technology and Rutgers University, Institute for Brain and Neuroscience Research, New Jersey Institute of Technology, Newark, NJ 07102, USA

**Author notes:** Corresponding Investigator, CONICET, Argentina.

## Abstract

Action potential generation in neurons depends on a membrane potential threshold, and therefore on how subthreshold inputs influence this voltage. In oscillatory networks, for example, many neuron types have been shown to produce membrane potential (*V_m_*) resonance: a maximum subthreshold response at a nonzero frequency. Resonance is usually measured by recording *V_m_* in response to a sinusoidal current (*I_app_*), applied at different frequencies (f), an experimental setting known as current clamp (I-clamp). Several recent studies, however, use the voltage clamp (V-clamp) method to control *V_m_* with a sinusoidal input at different frequencies (*V_app_*(*f*)) and measure the total membrane current (*I_m_*). The two methods obey systems of differential equations of different dimensionality and, while I-clamp provides a measure of electrical impedance (*Z*(*f*) = *V_m_*(*f*)/*I_app_*(*f*)), V-clamp measures admittance (*Y*(*f*) = *I_m_*(*f*)/*V_app_*(*f*)). We analyze the relationship between these two measurement techniques. We show that, despite different dimensionality, in linear systems the two measures are equivalent: *Z* = *Y*^−1^. However, nonlinear model neurons produce different values for *Z* and *Y*^−1^. In particular, nonlinearities in the voltage equation produce a much larger difference between these two quantities than those in equations of recovery variables that describe activation and inactivation kinetics. Neurons are inherently nonlinear and, notably, with ionic currents that amplify resonance, the voltage clamp technique severely underestimates the current clamp response. We demonstrate this difference experimentally using the PD neurons in the crab stomatogastric ganglion. These findings are instructive for researchers who explore cellular mechanisms of neuronal oscillations.

## 1 Introduction

Voltage and current clamp recording techniques are widely used in electrophysiological experiments to explore the properties of the ionic currents expressed in neurons and their functional effect in generating subthreshold and spiking activity [1–4]. Voltage clamp (V-clamp) experiments consist of measuring the current necessary to hold the voltage at a chosen level and involve a feedback amplifier. Current clamp (I-clamp) experiments, in contrast, consist on controlling the intensity of applied current and measuring the resulting changes in voltage and require no feedback loop. When the injected current is zero, I-clamp simply involves recording the intracellular voltage activity of neurons. I- and V-clamp experiments are complementary tools to investigate different aspects of neuronal dynamics. For instance, V-clamp is used to characterize the activation/inactivation curves and time constants of voltage-dependent ionic currents, while I-clamp is used to investigate dynamic properties of neurons such as the frequency-current relationships, sags exhibited by hyperpolarization and post-inhibitory rebound.

Dynamically, a primary difference between the I-clamp and V-clamp approaches is in the reduced dimensionality in V-clamp due to the elimination the time derivative of *V*, associated with the capacitive current. As a result, the V-clamp responses are typically less complex than the I-clamp ones. For example, for any constant value of *V*, the ionic current activation and inactivation dynamics are typically linear and one-dimensional, and therefore explicitly solvable. However, spontaneous activities, such as spiking and subthreshold voltage oscillations that depend on nonlinear mechanisms are only observable in I-clamp.

In spite of the reduced complexity of V-clamp as compared to I-clamp, both approaches have been used to investigate subthreshold (membrane potential) resonance, a description of preferred frequency responses of neurons to oscillatory inputs [5–54]. The presence of certain types of nonlinearities in the interaction among ionic currents (the current-balance equation) results in nonlinear amplifications of the voltage response to sinusoidal inputs as the input amplitude increases in I-clamp [55–57]. These nonlinear amplifications should be different in V-clamp, thus resulting in distinct resonance properties compared to those measured in I-clamp. However, for certain neuron types both approaches have shown to produce similar results [22].

Our goal is to compare the I-clamp and V-clamp responses of neurons to oscillatory inputs, in order to clarify conditions in which the two methods produce similar or distinct results. We use a variety of model neurons, ranging from linearized conductance-based models, to models with quadratic nonlinearities in the voltage equation, capturing the interaction between resonant currents (e.g., *I_h_* and the slow potassium current *I_M_*) and amplifying currents (e.g., the persistent sodium current *I_Nap_*) [56–58].

## 2 Methods

### 2.1 Models

#### 2.1.1 Linearized conductance-based models

We use linearized conductance-based models of the form

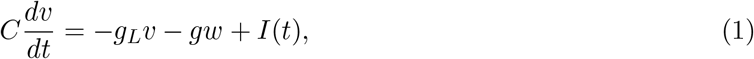

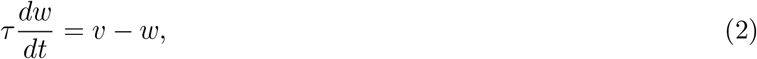

where *v* (mV) is voltage (with the resting potential translated to 0), *t* is time (ms),*g_L_* and *g* are linearized conductances (mS/cm^2^), C is the capacitance (*μ*F/cm^2^), *τ* is the linearized time constant (ms) and *I* is a time dependent current (*μ*A/cm^2^). The units of *w* are mV due to the linearization procedure used to obtain eqs. (1)–(2) from conductance-based models of Hodgkin-Huxley type [2]. We refer the reader to [6, 59] for details.

#### 2.1.2 Weakly nonlinear models

In order to capture some basic aspects of the differentiation between the nonlinear responses to oscillatory inputs in current and voltage clamp, we extend the linearized models to include simple types of nonlinearities with small coefficients either in the first or second equations.

The weakly nonlinear equations we use are

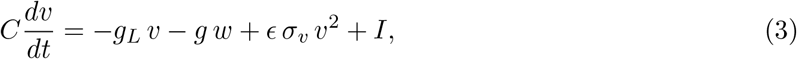

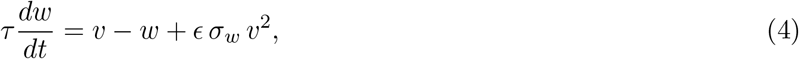

where *ϵ* is assumed to be small and the parameters 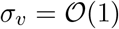 and 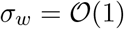. For simplicity we focus on nonlinearities that involve only the variable *v*.

#### 2.1.3 Caricature semilinear models

Generically, these models are of the form

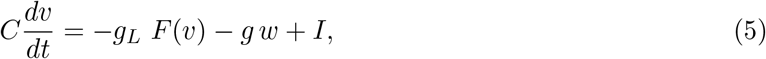

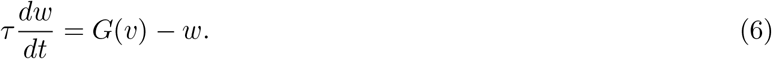

For *F* and *G* we use semi-sigmoidal nonlinearities of the form

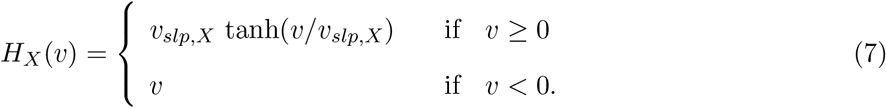

The function *H_X_*(*v*) (*X* = *v, w*) is continuously differentiable and semi-linear, with a sigmoid type of nonlinearity *v* ≥ 0, while it is linear for *v* < 0. Such functions allow one to investigate the effects of non-symmetric nonlinearities, on the model response to external inputs, as simple deformations of the linear nullclines. As we discuss later, they locally represent nonlinearities arising in neuronal models in the subthreshold voltage regime.

For *F*(*v*) = *G*(*v*) = *v*, system (5)–(6) reduces to the linear system (1)–(2). We refer to these models as LIN. We refer to the models with a nonlinearity in the first equation (*F*(*v*) = *H_v_* (*v*) and *G*(*v*) = *v*) as SIG-v and to these having a nonlinearity in the second equation (*F*(*v*) = *v* and *G*(*v*) = *H_w_*(*v*)) as SIG-w. Linearization of the SIG-v and SIG-w models yields LIN models with the same parameter values. In contrast to the more realistic quadratic models discussed below, the SIG-v and SIG-w models do not include an onset of spikes mechanism, and therefore allow for stronger input amplitudes than for the quadratic models.

#### 2.1.4 Quadratic conductance-based models

We will use quadratic models of the form

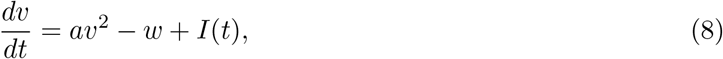

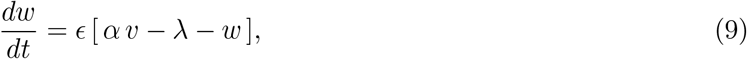

where *a, α, ϵ* and λ are constant parameters. These models can be derived from conductance-based models, with nonlinearities of parabolic type in the voltage equation, by using the so-called quadratization procedure described in [56, 60] (see also [61]). Examples of these models involve the interaction between *I_Nap_* and either *I_h_* or *I_M_* [56]. The units of the variables and parameters in eqs. (8)–(9) are [*v*] = mV, [*w*] = mV/ms, [*ϵ*] = 1/ms, [*a*] = 1/(ms mV), [*α*] = 1/ms, [λ] = mV/ms and [*I*] = mV/ms.

Note that Eqs. (3)–(4) can be thought of as a particular case of Eqs. (5)–(6) and are included in the quadratized formulation (8)–(9) after an appropriate change of variables.

#### 2.1.5 Oscillatory inputs: current (I-) and voltage (V-) clamp

In current and voltage clamp states, we use sinusoidal current and voltage inputs of the form

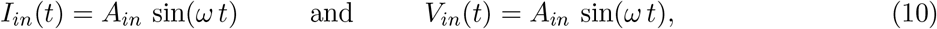

respectively, with

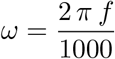

where *f* is the input frequency (Hz). Henceforth, we refer to the corresponding experiments as I-clamp and V-clamp respectively.

For input currents *I_in_*(*t*) the output is *V*(*t*) computed as the result of the corresponding system of differential equations. For voltage inputs *V_in_*(*t*) the output is *I*(*t*) computed by adding up all the other terms in the first equation (including the *dv/dt* term), after a proper rearrangement and using the variable w computed by using the remaining differential equation.

### 2.2 Voltage and current responses to sinusoidal inputs

#### 2.2.1 I-clamp: Impedance (*Z*-) and phase (Φ-) profiles

The voltage response *V_out_*(*t; f*) of a neuron to oscillatory current inputs *I_in_*(*t*) is captured by the neuron’s impedance, defined as the quotient between Fourier transforms of *V_out_* and *I_in_*:

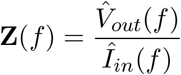

Impedance is a complex number with amplitude *Z*(*f*) and phase Φ(*f*). We refer to the impedance amplitude *Z*(*f*) simply as impedance and to the graphs *Z*(*f*) and Φ(*f*), respectively, as the impedance and phase profiles.

For linear systems receiving sinusoidal current inputs *I_in_*(*t*) as in eq. (10),

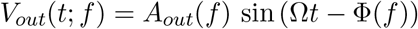

where Φ(*f*) is the phase shift between *I_in_* and *V_out_*, and

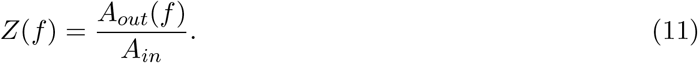

Linear systems exhibit resonance if *Z*(*f*) peaks at some non-zero (resonant) frequency *f_Z,res_* (Fig. 1-a1) and phasonance if Φ(*f*) vanishes at a non-zero (phasonant) frequency *f_Z,phas_* (Fig. 1-a2). For nonlinear systems, or linear systems with non-sinusoidal inputs, eq. (11) no longer provides an appropriate definition of impedance. Here we use the following definition

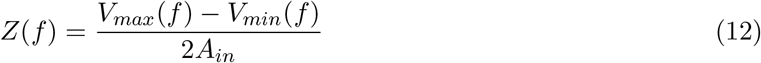

where *V_max_*(*f*) and *V_min_*(*f*) are the maximum and minimum of the steady-state oscillatory voltage response *V_out_*(*f*) for each value of f. For linear systems receiving sinusoidal inputs, eq. (12) is equivalent to eq. (11). Eq. (12) extends the concept of the linear impedance under certain assumptions (input and output frequencies coincide and the output amplitude is uniform across cycles for a given input with constant amplitude), which are satisfied by the systems we use in this paper. The resonant frequency *f_Z,res_* is then the peak frequency of *Z*(*f*) in eq. (12). Similarly to the linear case, the phase is computed as the distance between peaks of output and input normalized by period. We refer to the curves *V_max_*(*f*) and *V_min_*(*f*) as the upper and lower *Z*-envelopes, respectively (Fig. 1-a3).

**Figure 1:**
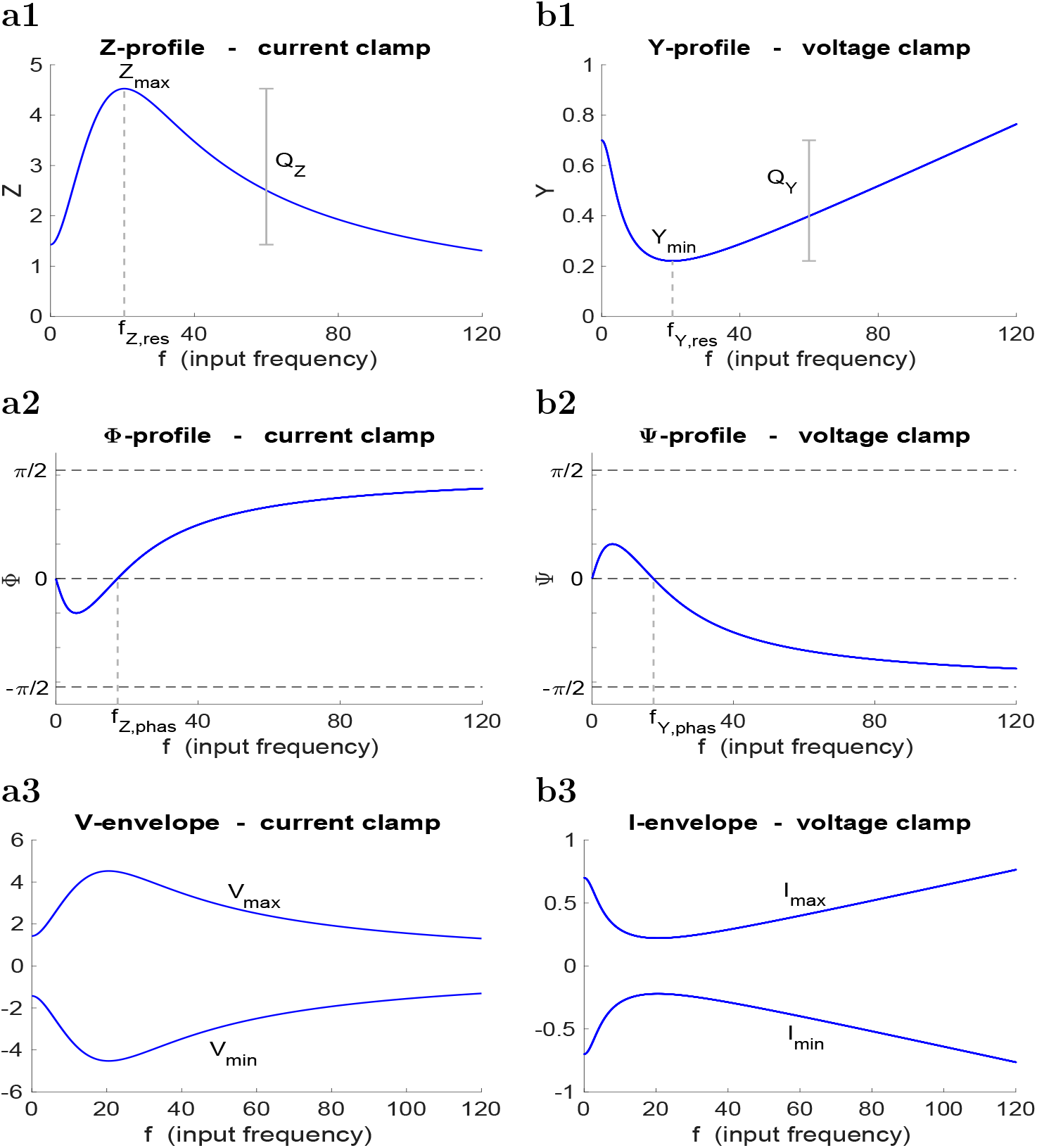
Schematic diagrams of the impedance and admittance amplitude and phase profiles. **(a1)** Impedance amplitude (*Z*) or, simply, impedance. The *Z*-resonant frequency *f_Z,res_* is the input frequency *f* at which the impedance *Z*(*f*) reaches its maximum *Z_max_*. The *Z*-resonance amplitude *Q_z_* = *Z_max_* – *Z*(0) measures the *Z*-resonance power. **(b1)** Admittance amplitude (*Y*) or, simply, admittance. The Y-resonant frequency *f_Y,res_* is the input frequency *f* at which the admittance *Y*(*f*) reaches its minimum *Y_min_*. The *Y*-resonance amplitude *Q_Y_* = *Y_min_* – *Y*(0) measures the *Y*-resonance power. **(a2)** Impedance phase (Φ) or, simple, Z-phase. The *Z*-phase-resonant frequency *f_Z,phas_* is the zero-crossing phase frequency. **(b2)** Admittance phase (Ψ) or, simply, Y-phase. The *Y*-phase-resonant frequency *f_Y,phas_* is the zero-crossing phase frequency. **(a3)** V-envelope. The upper (*V_max_*) and lower (*V_min_*) envelopes correspond to the maxima and minima of the voltage response for each input frequency. **(b3)** I-envelope. The upper (*I_max_*) and lower (*I_min_*) envelopes correspond to the maxima and minima of the current response for each input frequency.

#### 2.2.2 V-clamp: Admittance (*Y*) and phase (Ψ) profiles

The voltage response *I_out_*(*t; f*) of a neuron to oscillatory voltage inputs *V_in_*(*t*) is captured by the neuron’s admittance, defined as the quotient between the Fourier transforms of *I_out_* and *V_in_*:

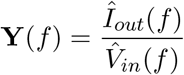

As with impedance, admittance is also a complex number, with amplitude *Y*(*f*) and phase Φ(*f*). As before, we refer to *Y*(*f*) simply as the admittance and to the graphs of *Y*(*f*) and Φ(*f*) as as the admittance and phase profiles, respectively.

For linear systems receiving sinusoidal current inputs *V_in_*(*t*) of the form (10),

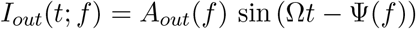

where Φ(*f*) is the phase difference between *V_in_* and *I_out_*, and

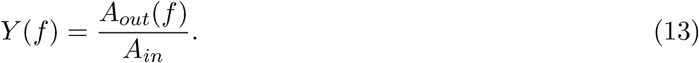

Linear systems exhibit resonance if *Y*(*f*) exhibits a trough at some non-zero (resonant) frequency *f_Y,res_* (Fig. 1-b1) and phasonance if Φ vanishes at a non-zero (phasonant) frequency *f_Y,phas_* (Fig. 1-b2). As with impedance, for nonlinear systems we use the following definition

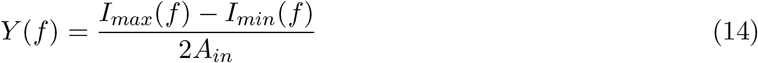

where *I_max_*(*f*) and *I_min_*(*f*) are the maximum and minimum of the oscillatory current response *I_out_*(*f*) for each value of *f*. For linear systems receiving sinusoidal inputs, eq. (14) is equivalent to eq. (13). The resonant frequency *f_Y,res_* is the frequency corresponding to the minimum of *Y* in eq. (14). Similarly to the linear case, the phase is computed as the distance between the peaks of the output and input normalized by the period. We refer to the curves *I_max_*(*f*) and *I_min_*(*f*) as the upper and lower I-envelopes, respectively (Fig. 1-b3).

#### 2.2.3 Inverse admittance (*Y*^−1^) and negative phase (−Ψ)

In general, the impedance (*Z*) and admittance (*Y*) are reciprocal quantities and can be used equivalently to characterize the response of a system to oscillatory inputs regardless of whether we are using I-clamp or V-clamp. Here, to avoid confusion, we use *Z* strictly for the voltage responses to current inputs and *Y* strictly for the current responses to voltage inputs. In order to compare the results for V-clamp and I-clamp, we will use the inverse admittance *Y*^−1^ and the negative phase −Ψ, measured in V-clamp, as comparable quantities to *Z* and Φ, measured in I-clamp. We refer to the corresponding curves as a function of the input frequency *f* as the inverse admittance (*Y*^−1^) and negative phase (−Ψ) profiles.

### 2.3 Experiments

Adult male crabs (*Cancer borealis*) were purchased from local seafood markets and kept in tanks filled with artificial sea water at 10-12 °C until use. Before dissection, crabs were placed on ice for 20-30 minutes to anesthetize them. The dissection was performed following standard protocols as described previously [62]. The STG was desheathed to expose the neurons for impalement. During the experiment, the preparation was superfused with normal *Cancer* saline (11 mM KCl, 440 mM NaCl, 13 mM CaCl_2_ · 2H_2_O, 26 mM MgCl_2_ · 6H_2_O, 11.2 mM Trizma base, 5.1 mM maleic acid; pH 7.4-7.5) at 10-12 °C. The PD neuron was identified by matching its intracellular activity with the extracellular action potentials on the lateral ventricular and PD motor nerves. Intracellular recordings were done using Axoclamp 2B amplifiers (Molecular Devices) with two intracellular electrodes, one for recording the membrane voltage and the second for current injection. Intracellular sharp glass electrodes were prepared using a FlamingBrown micropipette puller (P97; Sutter Instrument) and then filled with the electrode solution (0.6 M K_2_SO_4_ and 0.02 M KCl; electrode resistance 15-30 MΩ. To examine the response of the PD neuron in a range of frequencies, a chirp function *C*(*t*) was applied to the presynaptic neuron. This function can be described as:

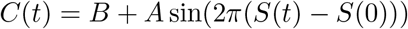

where *B* is the baseline, *A* is the amplitude and *S*(*t*) is a monotonically increasing function which determines the frequency range to be covered. When the chirp function was applied in voltage clamp, *B* = −45 mV and *A* = 15 mV. To obtain a larger sample set at the lower frequency range, we used a logarithmic chirp function by setting *S*(*t*) to be:

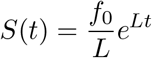

where

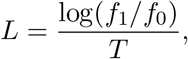

*f*_0_ (here 0.1 Hz) and *f*_1_ (here 4 Hz) are the initial and final frequencies in the chirp range, and *T* is its total chirp duration (here 100 s).

In both I-clamp and V-clamp experiments, the chirp functions were applied at least 3 times in control saline, and 3 times following bath application of 1 *μ*M proctolin (Bachem, USA). Impedance was measured using the fast Fourier transform function in MATLAB (MathWorks) as *FFT*(*V*(*t*)))/*FFT*(*I*(*t*)). Fit curves were done using a rational function with quadratic numerator and denominator.

## 3 Results

### 3.1 I-clamp and V-clamp produce different responses to the same constant inputs

Because of the reduced dimensionality of the system in V-clamp, as compared to I-clamp, the I-clamp response to constant inputs is expected to be more complex than its V-clamp response to a similar input. 2D linear systems, such as LIN: (1)-(2) in I-clamp, can display overshooting and damped oscillations that are absent in 1D linear systems, such as (1)-(2) in V-clamp (compare the blue curves in Figs. 2-b and 2-c1).

**Figure 2:**
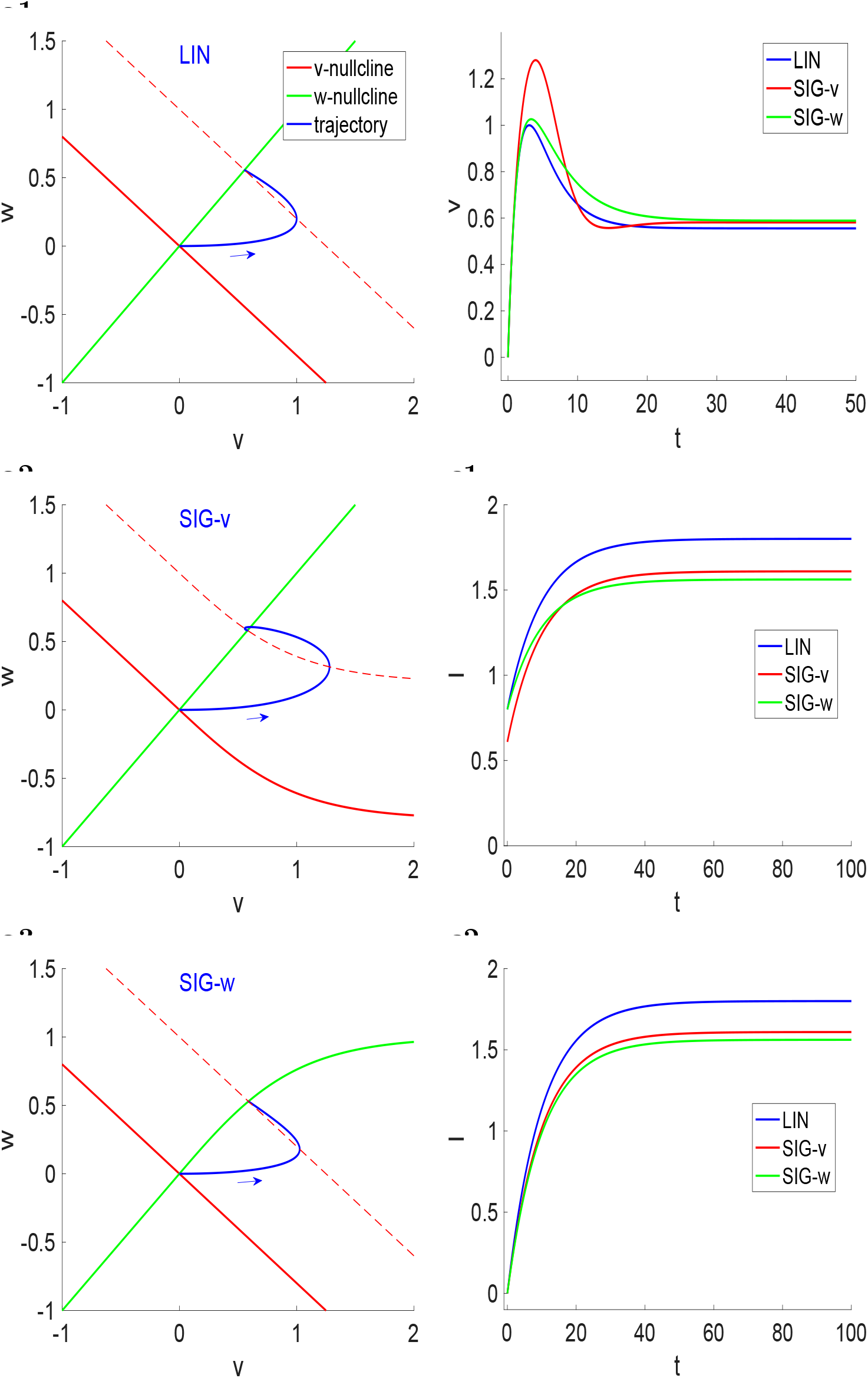
Dynamics of the autonomous LIN, SIG-v and SIG-w systems for representative parameter values. **(a)** Phase-plane diagrams (I-clamp) for the LIN (a1), SIG-v (a2), and SIG-w (a3) models for *I_app_* = 1. Trajectories are initially located at the fixed-point (0,0) for *I_app_* = 0. The dotted-red curve in each panel corresponds to the *v*-nullcline for *I_app_* = 1. **(b)** Voltage traces corresponding to panels a (current clamp). **(c)** Current traces (voltage clamp) for an applied voltage *V_app_* = 1. In panel c2 the initial conditions for the variable *w* were adapted so that the initial current is equal to zero. Parameter values: *C* =1, *g_L_* = 0.8, *g_L_* =1, *τ*_1_ = 10, *V_slp,v_* = 1, *v_slp,w_* = 1, *I_app_* = 1 (panels a and b), and *V_app_* = 1 (panel c)

In Figs. 2-a and 2-b we compare the I-clamp responses of the LIN, SIG-v and SIG-w models to constant inputs. The input and the common parameter values (*C, g_L_, g, τ*) are identical for all three models. We use the same values of *v_slp,v_* and *v_slp,w_* for the SIG-v and SIG-w models, respectively. The nonlinear *v*- (Fig. 2-a2) and w-(Fig. 2-a3) nullclines we use are, respectively, concave up for the SIG-v model (Fig. 2-a2) and concave down for the SIG-w model (Fig. 2-a3). This type of bending of the *v*- and *w*-nullclines are locally representative of the types of nonlinearities arising in neuronal models given the properties (i.e., monotonicity with respect to *v*) of the other nullcline. For monotonically increasing *w*-nullclines (as in Fig. 2-a2), the sigmoidal bending is the first stage in the generation of parabolic-like *v*-nullclines [58]. The use of the same type of nonlinearity for both the SIG-v and SIG-w models allows comparison of the dynamic effects produced by nonlinearities in the the two nullclines.

We define the response amplitude as the maximum value of *v* reached by the transient solution as it approaches the new steady state resulting from the constant input. In the phase-plane diagrams, this is determined by the intersection between the trajectory and the *v*-nullcline. In I-clamp, the SIG-v model exhibits a stronger nonlinear amplification of the response than the SIG-w or LIN model (Figs. 2-b). As seen in the phase-plane diagrams, because of the bending of the *v*-nullcline, the trajectory in panel a2 reaches further away from the fixed-point in the *v* direction than in panel a3 [55]. These differences become larger when the nonlinearities are more pronounced, i.e., for values of *v_slp,v_* and *v_slp,w_* (not shown).

In contrast, in the V-clamp responses there is little difference between the LIN, SIG-v and SIG-w models (Figs. 2-c). The dynamics of the w-equation in the LIN and SIG-v models are identical and *F*(*V_in_*) < *V_in_* for positive values of *V_in_* (eq. 7). Therefore, the curves *I*(*t*) are parallel and lower for the SIG-v model than for the LIN model (blue and red curves in Fig. 2-c1). The dynamics of the w-equation in the LIN and SIG-w models are different, but there are no nonlinear effects in the first equation. Since *G*(*V_in_*) < *V_in_* for positive values of *V_in_*, the fixed-point for the LIN model is higher than for the SIG-w model (blue and green curves in Fig. 2-c1). These relative relationships do not change if the initial conditions for *w* are not equal among the three models, but are chosen is such a way as to make the corresponding initial values of *I* equal (Fig. 2-c2).

### 3.2 I-clamp and V-clamp produce equivalent responses for linear systems receiving the same oscillatory input

In spite of the dimensionality differences between the I-clamp and V-clamp protocols, for linear systems and time-dependent inputs within a large enough class, both approaches produce the same response, so that *Z* = *Y*^−1^ and Φ = −Ψ (Fig. 3). We demonstrate this in detail for 3D linear systems and a complex exponential in Appendix A and provide the analytic solutions to generic 2D linear systems and linearized conductance based models in Appendix C. This result can be easily generalized to higher-dimensional linear systems and other types of time-dependent inputs by using their Fourier components.

**Figure 3:**
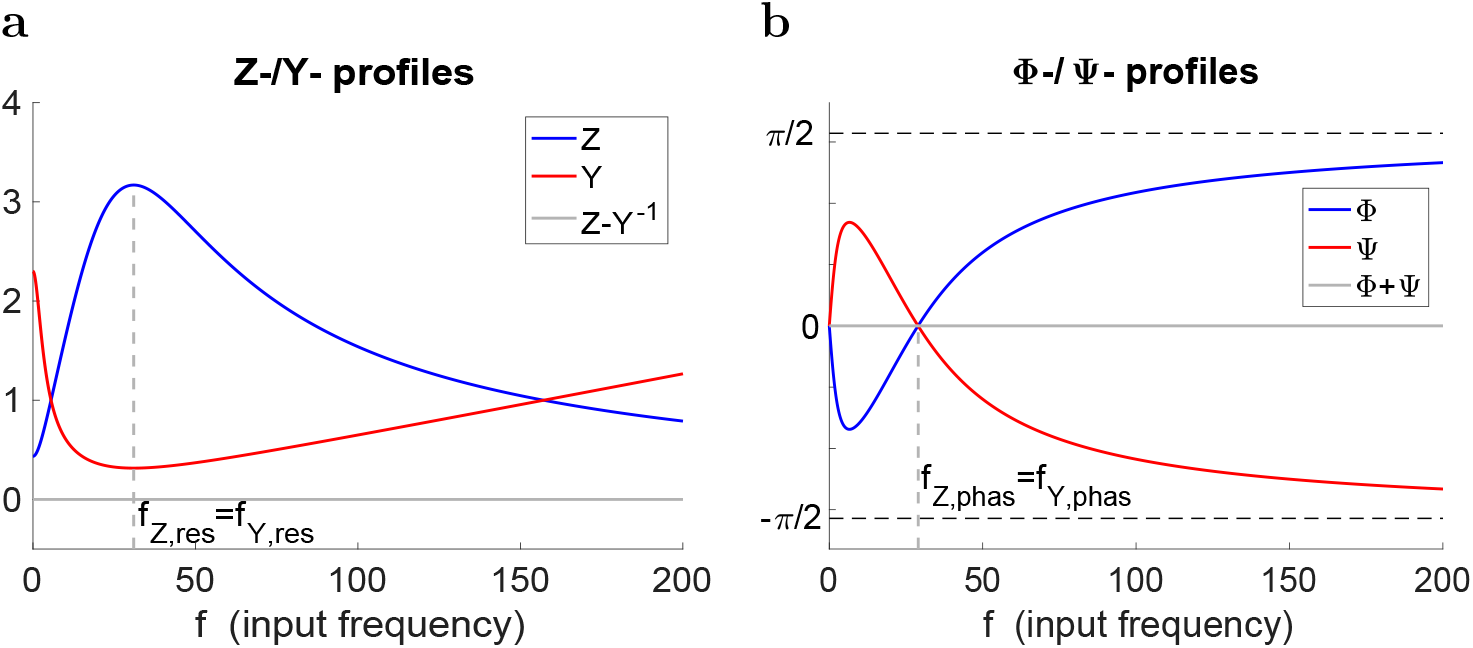
Equivalent impedance and admittance profiles for a representative linear system. **(a)** Impedance (*Z*) and admittance (*Y*) satisfying *Z* = *Y*^−1^ (gray line). **(b)** *Z*- and *Y*-phases (Φ and Ψ resp.) satisfying Φ = −Ψ (gray line). Parameter values: *C* =1, *g_L_* = 0.3, *g_L_* =2, *τ*_1_ =60 and *A_in_* = 1

The linear steady state responses to sinusoidal input satisfy three properties: (i) the input and output frequencies coincide, (ii) the input and output profiles are proportional, and (iii) the output envelope profiles for constant inputs are symmetric with respect to the equilibrium point from which they are perturbed. Note that (i) implies that the amplitude of the steady state response is uniform across cycles for each input frequency.

In terms of the I- and V-clamp responses to sinusoidal inputs, linearity implies that the *Z* (*Y*^−1^) and Φ (Ψ) profiles are independent of the input amplitude *A_in_*, and so are the *V* and *I*-envelope profiles when they are normalized by *A_in_* (see Fig. 1). A dependence of any of these quantities on *A_in_* indicates nonlinearity. Because of the symmetry property, the voltage and current envelopes (and the *Z* and *Y*^−1^ profiles) are redundant for linear systems [56].

The nonlinear models that we discuss in the next sections satisfy (i), but not necessarily (ii) and (iii). In previous work [55, 56] we have shown that for the quadratic model (8)-(9) and piecewise-linear models that capture the nonlinearities of the SIG-v and SIG-w models, increasing *A_in_* in I-clamp causes the impedance profile to increase in amplitude for input frequencies around the resonant frequency. We refer to this phenomenon as a nonlinear amplification of the voltage response. In the following sections we investigate how the nonlinearities in these models affect their response to current and voltage inputs and what similarities and differences arise between I-clamp and V-clamp.

### 3.3 Weakly nonlinear systems receiving oscillatory inputs: onset of the differences between I-clamp and V-clamp

We first consider weak nonlinearities in order to understand how differences between nonlinear *Z*- and *Y*^−1^- profiles emerge from the underlying linear profiles. We use the weakly nonlinear equations (3)–(4), where *ϵ* is assumed to be small and the parameters 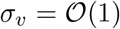 and 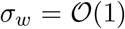. For *σ_v_* = *σ_w_* = 0 or, alternatively, *ϵ* = 0, eqs. (3)–(4) reduce to the linear model discussed in the previous section.

In the following, we will assume the capacitance parameter *C* is proportionally incorporated into the parameters *g_L_, g, I* and *σ_v_*. The time constant *τ*, in contrast, cannot be scaled away and its value affects the order of magnitude of the nonlinear term *ϵσ_w_v*^2^/*τ* in eq. 4. If 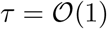, then this nonlinear term is 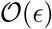. However, if 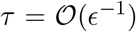, the nonlinear term is 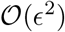 and therefore, to the 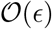 order, eq. 4 is linear and slow [55]. Finally, if 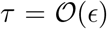, then the nonlinear term is 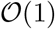, but, in this case, the other terms in eq. 4 are 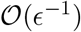 and therefore the negative feedback due to this equation is too fast for the underlying linear model to exhibit resonance.

We carry out a regular perturbation analysis for the system (3)-(4) in both I-clamp and V-clamp and we consider two scenarios for each: (i) a nonlinearity only in the v-equation (*σ_v_* = 1 and *σ_w_* = 0) or (ii) a nonlinearity only in the *w*-equation (*σ_v_* = 0 and *σ_w_* = 1). We refer to these models as WEAK-v and WEAK-w, respectively.

The 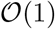 approximations (*ϵ* = 0) in both cases (I-clamp and V-clamp) are the linear systems discussed in the previous section for which *Z*(*ω*) = *Y*^−1^(*ω*) and Φ(*ω*) = −Ψ(*ω*). We show that the 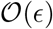 approximation systems are different for I-clamp and V-clamp. In particular, with all other parameters equal, (i) a nonlinearity in the *v*-equation (3) has a stronger effect on the *Z* and *Y*^−1^ profiles than the same nonlinearity in the *w*-equation (4), and (ii) the effect of the nonlinearity is stronger on the *Z* profile than on the *Y*^−1^ profile. As a result, the 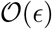 components and therefore the overall impedance and admittance are also different.

The results presented below for system (3)-(4) are based on the results for general 2D systems with weak nonlinearities presented in detail in the Appendix D. Along this section, we frequently refer to these results and the solutions for generic 2D linear systems presented in the Appendix C.

#### 3.3.1 Asymptotic approximation in I-clamp

We expand the solutions of (3)-(4) with *I* = *A_in_* sin(*ω t*) in series of *ϵ*

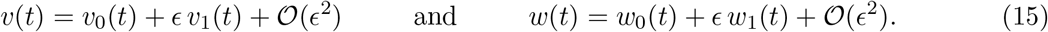

Substituting into (3)–(4) and collecting the terms with the same powers of *ϵ* we obtain the following systems for the 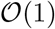 and 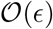 orders, respectively.

##### 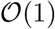 system

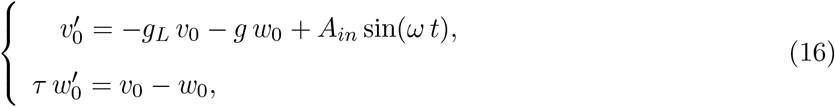

The solution to this linear system (Appendix C.3) is given by

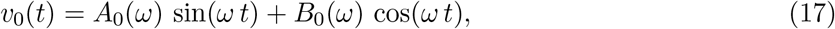

where *A*_0_(*ω*) and *B*_0_(*ω*) are given by (74), and

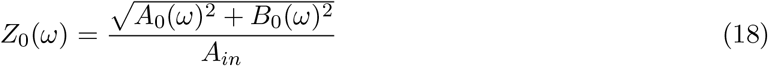

is the impedance for the linear system.

##### 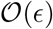 system

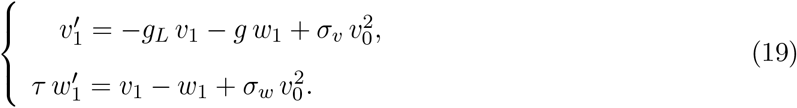

The solution to this forced linear system (Appendix D.1) is given by

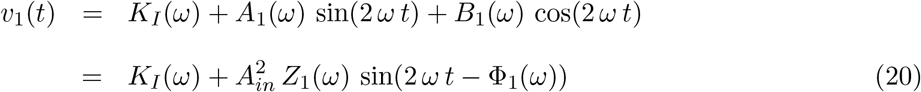

where

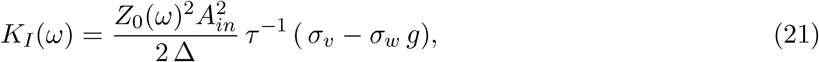

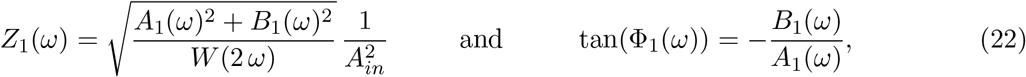

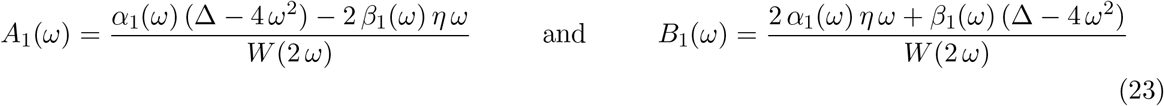

with Δ and *η* given by (75), and *W*(2 *ω*) given by (57) with the frequency multiplier *k* = 2,

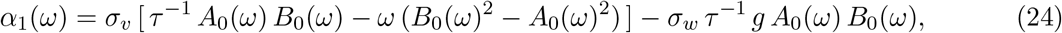

and

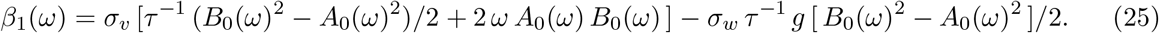

Note that *K_I_* and *Z*_1_(*ω*) are independent of *A_in_*.

#### 3.3.2 The onset of nonlinear effects in I-clamp: The nonlinear effects are stronger for the WEAK-v model than for the WEAK-w model

To the 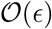 order of approximation,

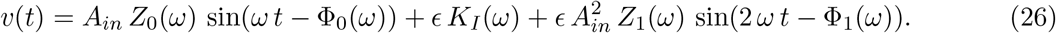

The frequency-dependent constant term *K_I_*(*ω*) affects the *v*-envelope (Fig. 4-a1) in a frequency-dependent manner, but has at most a negligible effect on *Z*, which involves the difference between the upper and lower envelopes and not the envelopes themselves. (Note that the difference between the upper and lower envelope is zero at the 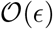 order.) *K_I_*(*ω*) depends on *τ* only through the dependence of *Z*_0_(*ω*) on *τ*, since the explicit occurrence of the *τ* term is canceled out by its occurrence in the denominator of Δ in eq. (75). Therefore *K_I_*(*ω*) has an effect on the v-envelope even for large values of *τ*.

**Figure 4:**
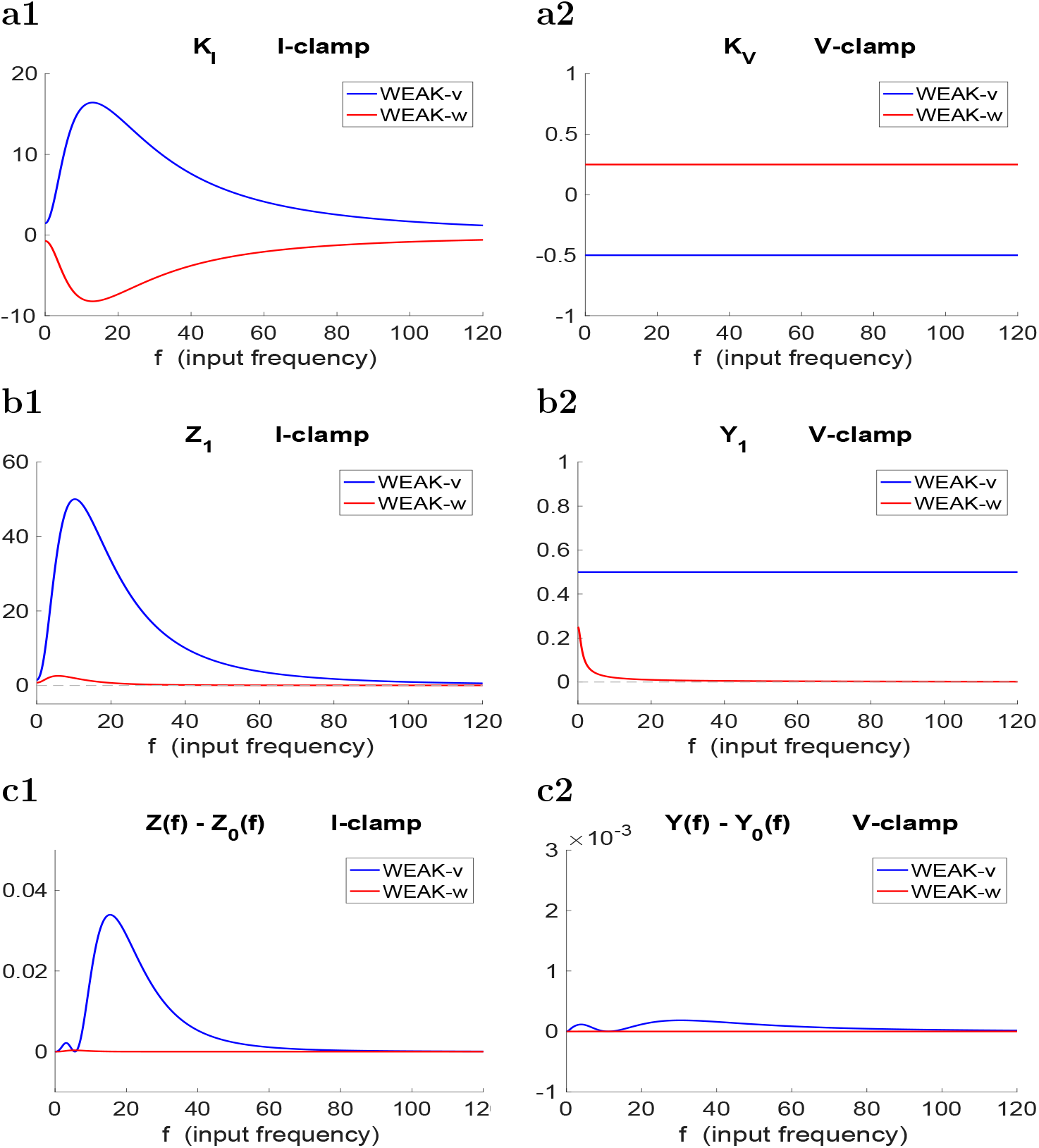
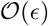 approximation to the voltage and current responses to sinusoidal inputs for the weakly nonlinear WEAK-v and WEAK-w models (3)-(4). **(a)** Constant terms in the 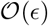 approximation of *Z* and *Y*. **(a1)** *K_I_* in (20). **(a2)** *K_V_* in (34). **(b)** Coefficients of the oscillatory terms in the 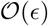 approximation of *Z* and *Y*. **(b1)** *Z*_1_ in (20). **(b2)** *Y*_1_ in (34). **(c)** Difference between *Z* and *Y* for the weakly nonlinear models and the 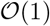 approximations (linear system). **(c1)** *Z*(*f*) – *Z*_0_(*f*). **(c2)** *Y*(*f*) – *Y*_0_(*f*). The *Z*- and *Y*-profiles for the WEAK-v and WEAK-w models up to the 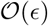 terms was calculated using (12) and (14), respectively. Parameter values: *C* =1, *g_L_* = 0.2, *g_L_* = 0.5, *τ*_1_ = 100, *A_in_* = 1 and *τ* = 0.01. For the WEAK-v model we used *σ_v_* = 1 and *σ_w_* =0 and for the WEAK-w model we used *σ_v_* =0 and *σ_w_* = 1.

From (21) follows that *K_I_*(*ω*) is positive for the WEAK-v model and negative for the WEAK-w model (Fig. 4-a1). Had this been the only term in the 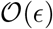 correction for *v*, the nonlinear effects would have been amplified in the WEAK-v model and attenuated in the WEAK-w model. The origin of *K_I_*(*ω*) is the quadratic form of the nonlinearity through the trigonometric transformation of squared sinusoidal functions into into standard sinusoidal functions. Constant terms will not necessarily be present for other types of nonlinearities (e.g., cubic).

The effects of *Z*_1_(*ω*) are more difficult to analyze due to its complexity. It is instructive to examine the effects of *Z*_1_(*ω*) in the limit of large values of *τ* where eq. (22) is simplified. We show in the Appendix E.1 that *A*_0_(*ω*) and *B*_0_(*ω*) in eq. (18) are 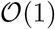 quantities for large enough values of *τ*. Therefore, for large enough *τ*, the terms that explicitly depend on *τ* in both *α*_1_ (*ω*) and *β*_1_(*ω*) are 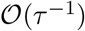, while the remaining terms are 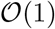. In particular, the terms involving *σ_w_* are 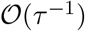. This implies that for large enough *τ*, the nonlinearity in the WEAK-v model may have a relatively strong effect on the impedance and phase profiles, while the nonlinearity in the WEAK-w model may have a much weaker effect. This difference persists for smaller values of *τ* away from the limit (Fig. 4-b1).

It is important to note that *Z*_1_(*ω*) is not the 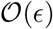 correction to the *Z*(*ω*)-profile. This correction is *Z*(*ω*) – *Z*_0_(*ω*) and is affected not only by *K_I_*(*ω*) and *Z*_1_(*ω*), but also by the fact that the sinusoidal term involves frequencies twice as large as the frequency of *Z*_0_(*ω*), as expected from the presence of quadratic nonlinearities. Fig. 4-c1 shows that the 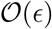 correction to *Z*(*ω*) is more pronounced for the WEAK-v model than for the WEAK-w model and the nonlinear amplification of *Z*(*ω*) peaks at the resonant frequency band.

An additional point to note is the effect of the presence of 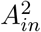 in the 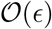 correction to *v*. For values of *A_in_* < 1, the effects of *A_in_* will be attenuated, while for values of *A_in_* > 1 they will be amplified. This effect is the same for both models.

#### 3.3.3 Asymptotic approximation in V-clamp

We expand the solutions of (3)-(4) with *v* = *A_in_* sin(*ωt*) in series of *τ*:

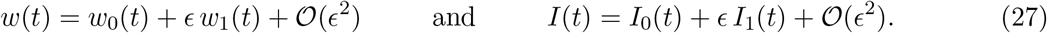

Substituting into (3)-(4) and collecting the terms with the same powers of *ϴ* we obtain the following systems for the 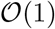 and 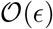 orders, respectively.

##### 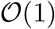 system

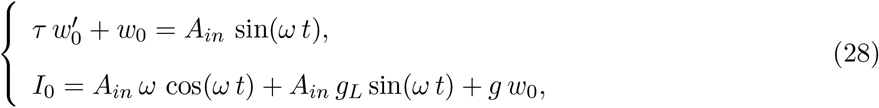

The solution to the first equation in (28) is given by (77) in Appendix C.4. Substitution into the second equation in (28) yields

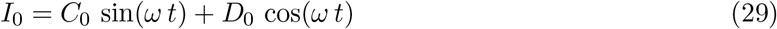

where *C*_0_(*ω*) and *D*_0_(*ω*) are given by (79), and

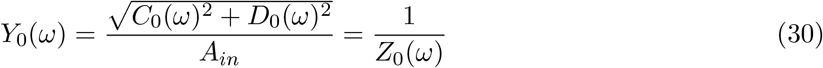

is the admittance for the linear system.

##### 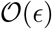 system

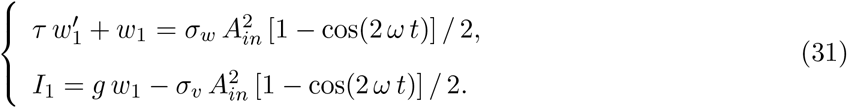

The solution (Appendix D.2) is given by

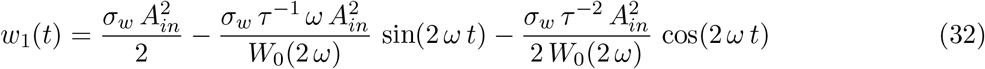

with *W*_0_(2 *ω*) given by (62) with *k* = 2 and *d* = −*τ*^−1^. Substitution into the second equation in (97) yields

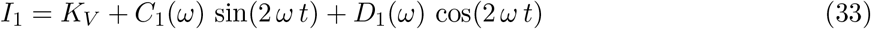

or

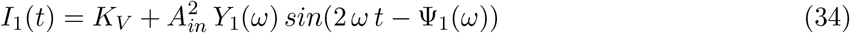

where

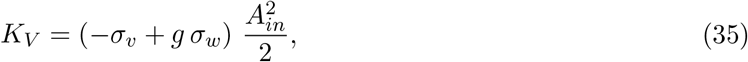

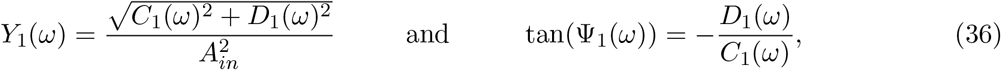

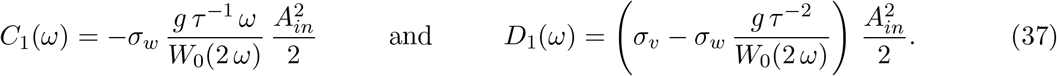

#### 3.3.4 The onset of nonlinear effects in V-clamp: The nonlinear effects are stronger for the WEAK-v model than for the WEAK-w model

To the 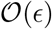 order of approximation

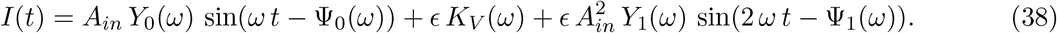

Similarly to the I-clamp protocol discussed above, the term *K_V_* (*ω*) affects the I-envelope (Fig. 4-a2) in a way that has at most a negligible effect on *Y*, but in contrast to the I-clamp protocol, this term is independent of both *ω* and *τ*. As for I-clamp protocol, *K_V_* originates in the quadratic nonlinearity and may not be present in other types of nonlinearities.

From (35) follows that *K_V_* is negative for the WEAK-v model and positive for the WEAK-w model (Fig. 4-a2). Had this been the only term in the 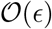 correction for I, the nonlinear effects would have been amplified in the WEAK-v model and attenuated in the WEAK-w model in terms of the inverse impedance *Y*^−1^. However, in Appendix E.2 we show that, for large enough values of 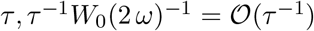 and 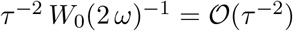 (in the limit *τ* → ∞ both quantities approach zero). Therefore, for large values of *τ* both *C*_1_(*ω*) and the second term in *D*_1_(*ω*) are negligible. The remaining term in *D*_1_(*ω*) is *σ_v_ A_in_*/2, which is independent of both *τ* and *ω*. As with the I-clamp protocol, this implies that, in V-clamp, for larger enough values of *τ* the nonlinearity in the WEAK-v model may have a relatively strong effect on the admittance and phase profiles, while the nonlinearity in the WEAK-w model will have a much weaker effect (Fig. 4-b2).

In Fig. 4-b3 we show the 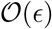 correction to the *Y*(*ω*)-profile, which is affected by *K_V_*(*ω*), *Y*_1_(*ω*), and the sinusoidal term involves frequencies twice as large as the frequency of *Y*_0_(*ω*). This 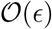 correction is more pronounced for the WEAK-v model than for the WEAK-w model.

#### 3.3.5 The 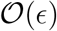 correction to the *Z*-profile is larger than the 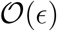 correction to the *Y*-profile and these nonlinear differences increase with increasing values of *A_in_*

We first examine this for large values of *τ* where, as we showed above, all the involved expressions are simplified. From Appendices E.3 and E.2, for large enough values of *τ*

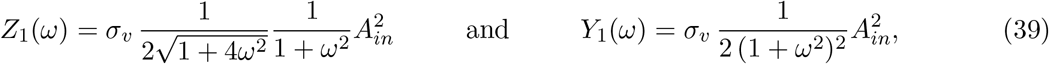

respectively. Therefore

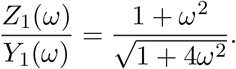

This quotient is equal to 1 for *ω* = 0 and is an increasing function of *ω*, indicating that the frequency-dependent 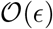 correction to *Z*_0_(*ω*) is larger than the 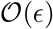 correction to *Y*_0_(*ω*). This behavior is illustrated in Figs. 4-b1 and -b2 for *τ* = 100. Comparing the corresponding blue curves (WEAK-v model) and red curves (WEAK-w model) shows that, for each model, *Z*_1_ > *Y*_1_ and the difference is much larger for *Z*_1_ than for *Y*_1_. Figs. 4-c1 and -c2 show that the 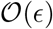 approximation to the *Z*-profile is larger than the 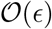 approximation to the *Y*-profile and, in both cases, the 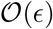 approximations to the *Z*- and *Y*-profiles are much stronger for the WEAK-v than for the WEAK-w models. A similar behavior occurs for the frequency-independent 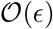 corrections. Comparing Figs. 4-c1 and -c2 illustrates that the 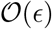 correction to the *Z*-profile is more pronounced than the 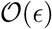 correction to the *Y*-profile.

### 3.4 I-clamp and V-clamp responses for the SIG-v and SIG-w models

Here and in the next section we investigate the effects of the model nonlinearities on the I-clamp and V-clamp responses to oscillatory inputs in the SIG-v and SIG-w models (5)-(6) for which we cannot find an analytical approximate solution. These nonlinear caricature models are useful because they are natural extensions of the linearized models discussed above and, unlike the more realistic quadratic model discussed below, lack parameter regimes where the solutions increase without bound, thus allowing for the use larger input amplitudes.

#### I-clamp and V-clamp produce nonlinear amplifications of the *Z*- and *Y*^−1^-profiles in the SIG-v, but not the SIG-w model

Figs. 5-a1 and -b1 show that for the SIG-v model both the *Z*- and *Y*^−1^-profiles (red) are larger than the corresponding linear ones (blue). In addition to the nonlinear amplification, the v-response for the SIG-v model peaks at a lower frequency than the LIN model. In contrast to the SIG-v model, the *Z*- and *Y*^−1^-profiles (green) are not nonlinearly amplified (panels a1 and b1). However, *f_res,Z_* (panel a1) is lower than the linear prediction (blue). The nonlinear amplification of *Z*- and *Y*^−1^-profiles for the SIG-v model is more pronounced the stronger the nonlinearities (the smaller the values of *v_s1p,v_* and *v_s1p,w_*), but the SIG-w model still exhibits quasi-linear behavior in these cases (not shown). Consistent with previous work [55], the nonlinear amplification of the *Z*-profile for the SIG-v models is less pronounced for smaller values of *τ* (due to smaller time scale separation between *v* and *w*; not shown).

**Figure 5:**
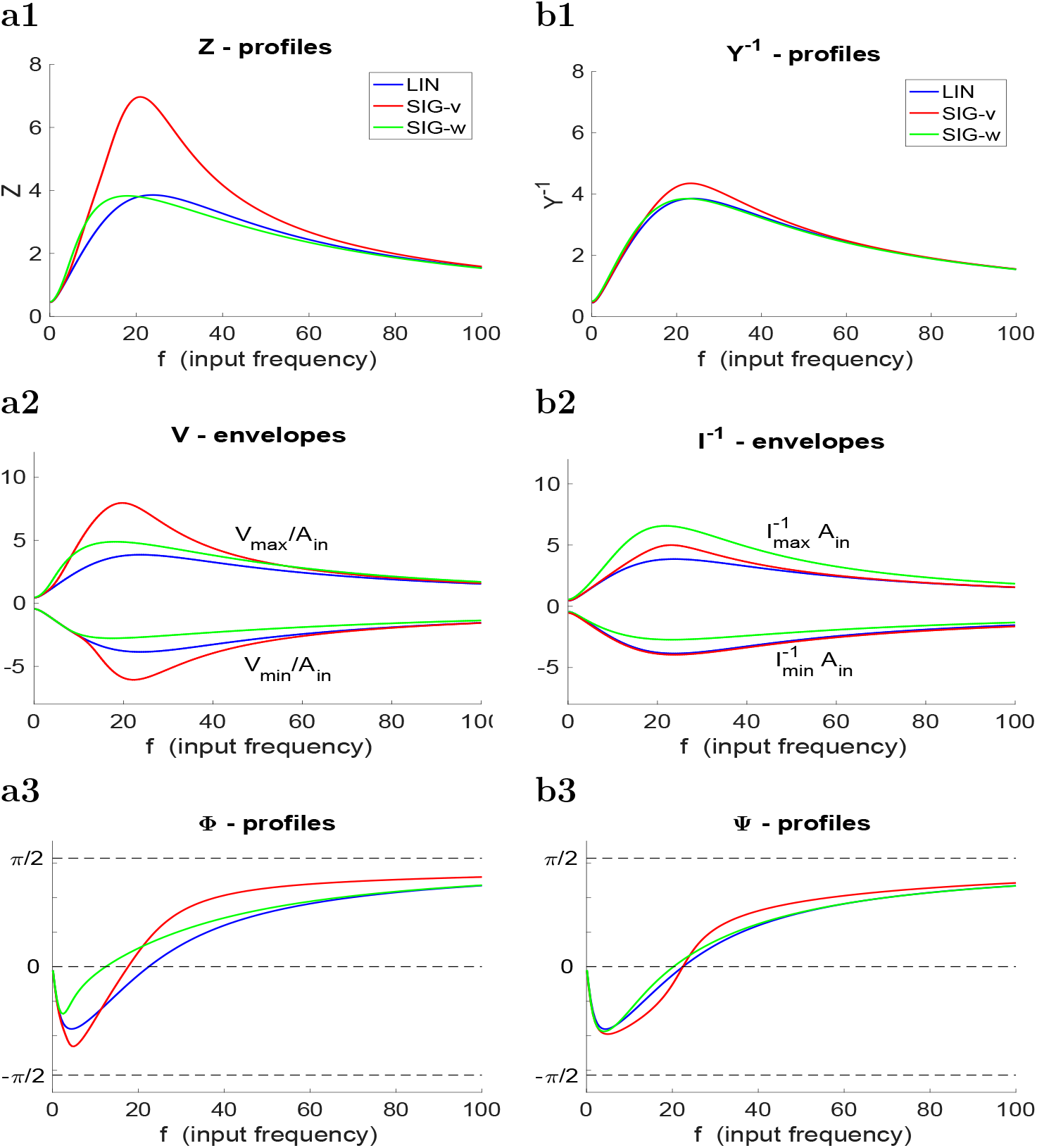
Linear and nonlinear voltage and current responses to sinusoidal current and voltage inputs respectively for a representative set of parameter values. **(a)** Voltage response to sinusoidal currents inputs. **(b)** Current response to sinusoidal voltage inputs. Parameter values: *C* =1, *g_L_* = 0.25, *g* = 2, *τ*_1_ = 100, *V_sip,sgv_* = 1, *V_sip,sgw_* = 1, *I_in_* = 1 (panels a) and *V_in_* = 1 (panels b).

#### I-clamp and V-clamp produce nonlinear changes in the *V*- and *I*^−1^-envelopes in both the SIG-v and SIG-w models

The behavior of the *V*- and *I*^−1^-envelopes (Figs. 5-a2 and -b2) for the SIG-w model is different from the SIG-v and LIN models. Although the *Z*- and *Y*^−1^-profiles show quasi-linear behavior, both the upper and lower envelopes are displaced above the linear ones (green). In other words, the upper envelopes are nonlinearly amplified, while the lower envelopes are nonlinearly attenuated. Because the *V*- and *I*^−1^-envelopes are displaced almost in a parallel manner, the *Z*- and *Y*^−1^-profiles remain almost identical to the linear ones. This implies that the *Z*- and *Y*^−1^ profiles for the SIG-w model fail to capture significant nonlinear aspects of the corresponding *V*- and *I*^−1^ envelopes, and therefore are not good predictors of the voltage and current response behaviors. This observation is important, particularly when one wants to infer the neuronal suprathreshold resonant properties from the subthreshold ones, where the action potential threshold depends primarily on the upper *V*-envelope. The differences between the SIG-v and SIG-w models are more pronounced for stronger nonlinearities, but the lower *I*^−1^-envelope for the SIG-v model remains almost unaffected by changes in *A_in_* (not shown).

#### The nonlinear amplification of the response for the SIG-v model is significantly stronger for I-clamp and V-clamp

Comparing Figs. 5-a1 and -b1 shows that the differences in the *Z*-profiles for the SIG-v and LIN models (red and blue in panels a1 and b1) are significantly larger than the differences in the *Y*^−1^ profiles for the same models. The same type of behavior is observed in the corresponding *V*- and *I*^−1^-envelopes (panels a2 and b2). These phenomena are stronger with more pronounced nonlinearities, but the relative differences between *Z* and *Y*^−1^ persist (not shown). The nonlinear amplification *Y*^−1^-profile for the SIG-v model is less affected by decreasing values of *τ* (not shown). Note that changes in *τ* change both *f_Z,res_* and *f_Y,res_* and therefore the *Z*- and *Y*^−1^ profiles are displaced with respect to the values in Figs. 5-a1 and -b1.

#### I-clamp captures nonlinear phase-shift effects in both the SIG-v and SIG-w models better than V-clamp

The Φ-profiles in Fig. 5-a3 show that *f_Z,phas_* both both the SIG-v and SIG-w models (red and green) are not well approximated by *f_Z,phas_* for the LIN model, with *f_Z,phas_* for the SIG-w model smaller than for the SIG-v model. In contrast, *f_Y,phas_* for the SIG-v and LIN models are almost equal and slightly higher than *f_Y,phas_* for the SIG-w model. This behavior persists for stronger nonlinearities, although the differences increase as the nonlinearities become more pronounced (not shown). This behavior also persists for other values of *τ* although both *f_Z,phas_* and *f_Y,phas_* change with *τ* (not shown).

### 3.5 I-clamp and V-clamp responses for the quadratic model

We now extend our investigation of the I-clamp and V-clamp responses to oscillatory inputs to the more realistic parabolic *v*-nullcline in the quadratic model (8)-(9) (Fig. 6). Such quadratic models can be derived from conductance-based models that describe the interaction between a regenerative current (e.g., persistent sodium) and a restorative current (e.g., h- or M-), when the voltage (*v*-) nullcline in the subthreshold voltage regime is parabolic [56, 60].

**Figure 6:**
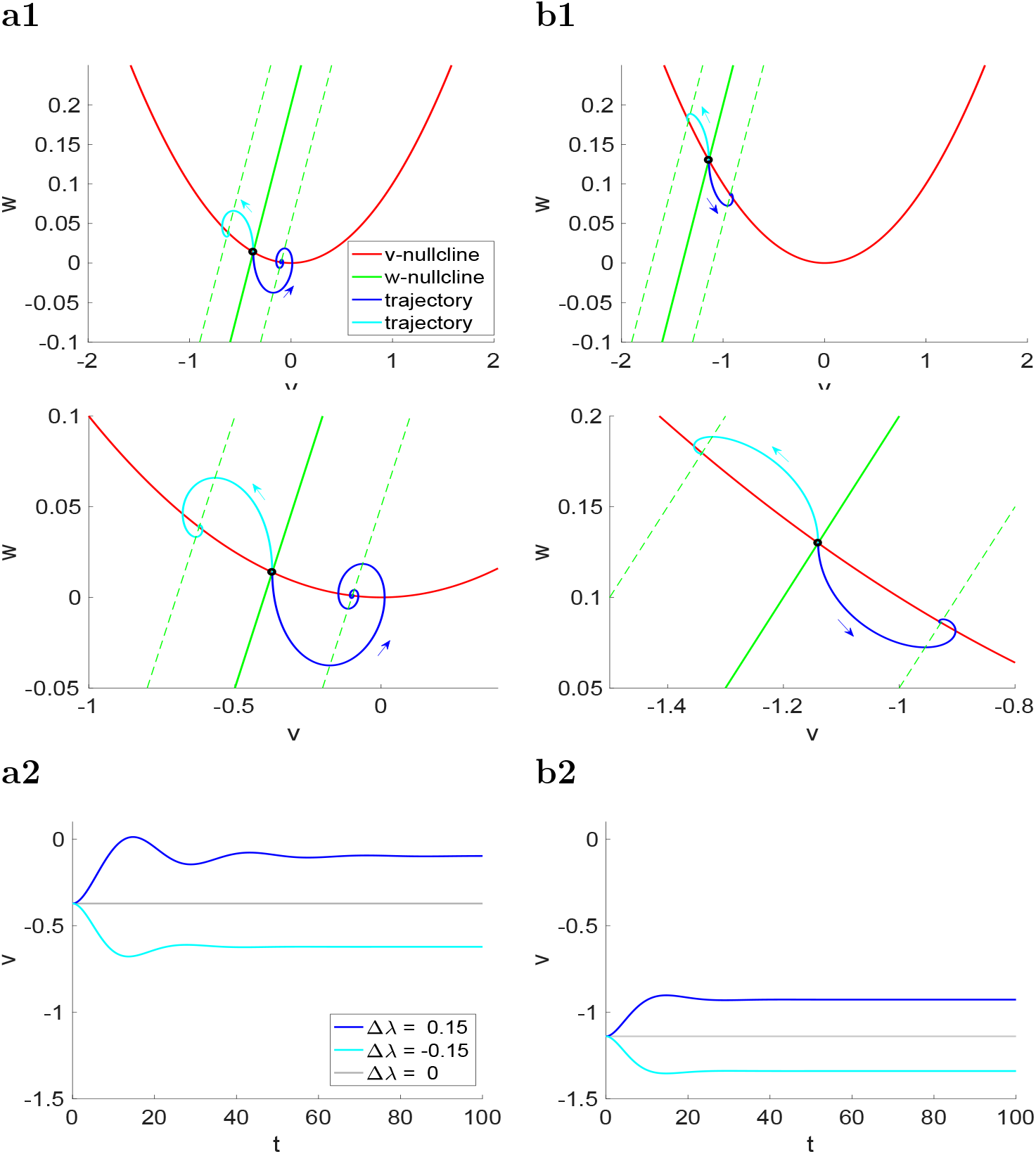
Dynamics of the autonomous quadratic model for representative parameter values and levels of λ. **(a)** λ = −0.2. **(b)** λ = −0.7. λ represents the baseline applied (DC) current, which determines the steady state values of *v* and the fixed-point (black dot on the intersection between the nullclines in panels a). The parameter Δλ represents constant deviations from λ. **Panels a1 and b1**. Superimposed phase-plane diagrams for Δλ = 0, Δλ = 0.15 and Δλ = −0.15. The solid-green w-nullcline corresponds to Δλ = 0 and the dashed-green *w*-nullclines correspond to Δλ = ±0.15. Their intersection with the red *v*-nullcline determines the fixed-points for the perturbed systems. Trajectories initially at the fixed-point for Δλ = 0 evolve towards the perturbed fixed-points for Δλ = 0.15 (blue) and Δλ = −0.15 (cyan). The closer the fixed-points to the knee of the *V*-nullcline, the more nonlinear and amplified is the response. The bottom panels are magnifications of the top ones. **Panels a2 and b2**. Voltage traces. Parameter values *a* = 0.1, *α* = 0.5, *ϵ* = 0.1.

When the resting potential (fixed-point) is away from the knee of the *v*-nullcline (Fig. 6-b1) the dynamics are quasi-linear, as reflected in the symmetry of the system’s response (Fig. 6-b2) to positive (blue curve) and negative (cyan curve) constant inputs relative to the resting potential (gray). When the resting potential is closer to the minimum of the *v*-nullcline (Fig. 6-a1), this symmetry is broken due to the presence of the parabolic nonlinearity and responses to positive constant inputs are more amplified than responses to negative inputs (Figs. 6-a1 and -a2).

Fig. 7 shows that the principles extracted from the previous models regarding the differences between I-clamp and V-clamp persist for the quadratic model (8)-(9). Panels a and b correspond to the same model parameters except for *ϵ* (corresponding to the value of *τ* in the weakly nonlinear models investigated above), which is larger in panels a (*ϵ* = 0.01) than in panels b (*ϵ* = 0.05. The values of *A_in_* in each case were adjusted to be just below the threshold value for which the solution increases without bound, representing the generation of spikes.

**Figure 7:**
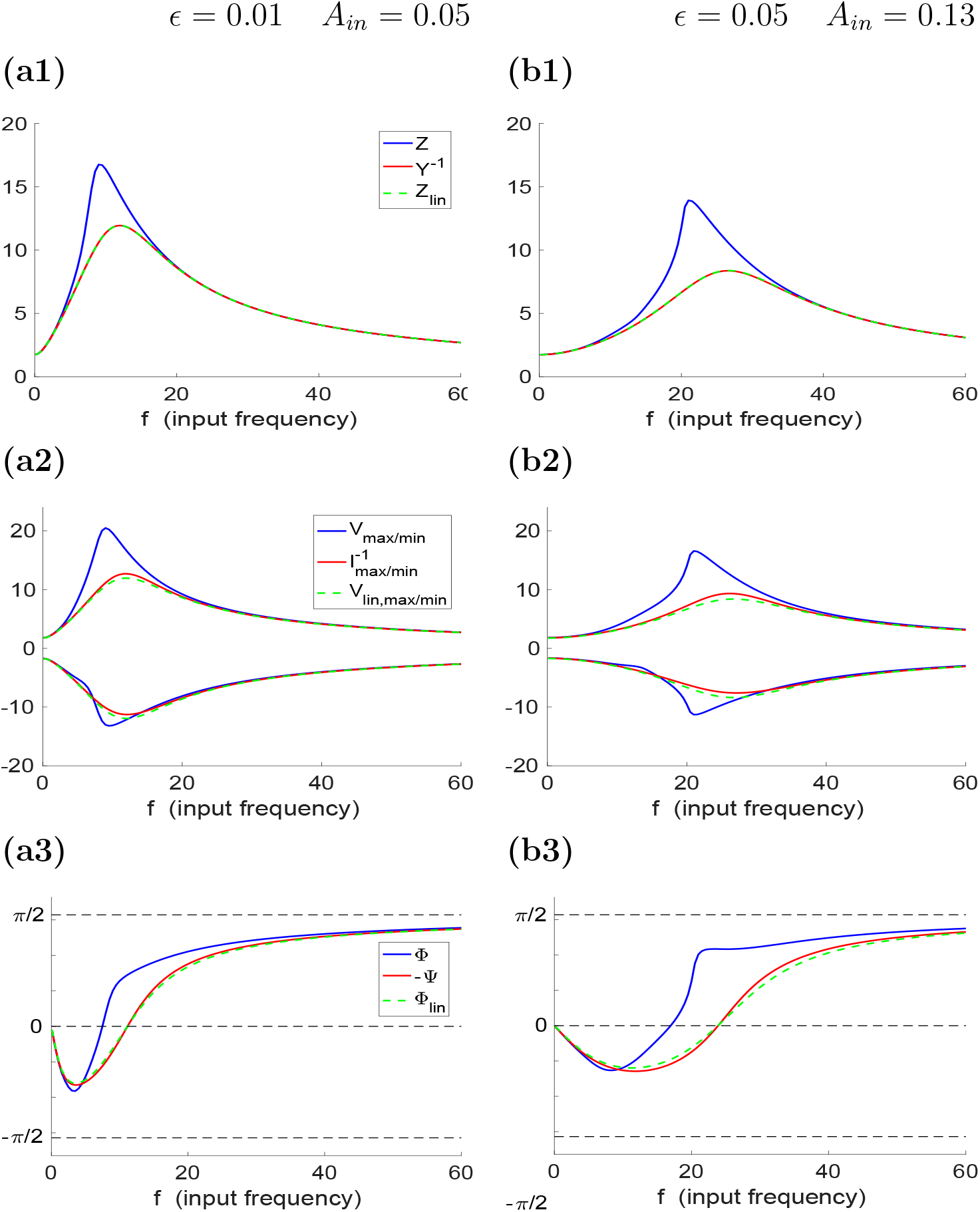
Voltage and current responses to sinusoidal current and voltage inputs respectively for the quadratic and linearized quadratic models for representative parameter values. **(a)** *ϵ* = 0.01 and *A_in_* = 0.05. **(b)** *ϵ* = 0.05 and *A_in_* = 0.13. The value of *A_in_* in both cases is below, but close to the threshold value for spike generation. **Top panels**. *Z*- and *Y*^−1^-profiles. Middle panels. *V* and *I*^−1^ envelopes. **Bottom panels**. Φ- and Ψ-profiles. Parameter values *a* = 0.1, *α* = 0.5, λ = −0.2.

In both cases, V-clamp responses (*Y*^−1^, red) almost coincide with linear responses (*Z_lin_*, green), while I-clamp responses (*Z*, blue) are nonlinearly amplified. The red voltage envelopes (Fig. 7-a2 and -b2) are displaced with respect to the green ones due primarily to the constant terms generated by the quadratic nonlinearities. This has almost no effect on the corresponding *Z_lin_*- and *Y*^−1^-profiles (Fig. 7-a1 and -b1). The amplification of the blue voltage envelopes is not symmetric, reflecting the presence of quadratic nonlinearities [56]. Because of the stronger time scale separation the *Z*- and *Y*^−1^-profiles for *ϵ* = 0.01 (Fig. 7-a1) shows a sharper peak compared to the *Z*- and *Y*^−1^-profiles for *ϵ* = 0.05.

### 3.6 I-clamp and V-clamp responses in biological neurons

To examine the predictions of our mathematical analysis, we explored measurements of impedance and admittance in the pyloric dilator (PD) neurons of the crab stomatogastric ganglion. PD neurons have been shown to produce resonance and their impedance profiles have been measured in both I-clamp and V-clamp [62–64]. Additionally, it is known that the modulatory neuropeptide proctolin activates a modulatory-activated inward current (*I_MI_*) in these neurons. The voltage-dependence of *I_MI_* and its kinetics are very similar to that of the persistent sodium current *I_Nap_* [65]. We therefore predicted that proctolin should amplify the resonance properties [5] of the PD neuron, and sought to measure the effect of proctolin in both V-clamp and I-clamp conditions.

A comparison between these two cases is shown in Fig. 8. We measured the response of the neuron in I-clamp (Fig. 8a1) and V-clamp (Fig. 8a2), respectively, by injecting a sweeping-frequency sinusoidal chirp function as a current or voltage input. Because the PD neuron resonance frequency is close to 1 Hz, we allowed the chirp function to sweep frequencies from 0.1-4 Hz. As predicted from our analysis, in I-clamp, addition of proctolin greatly enhanced the peak of the *Z*-profile (Fig. 8b1), whereas, in V-clamp the same modulatory effect only produced a moderate change in the *Y*^−1^-profile (Fig. 8b2).

**Figure 8:**
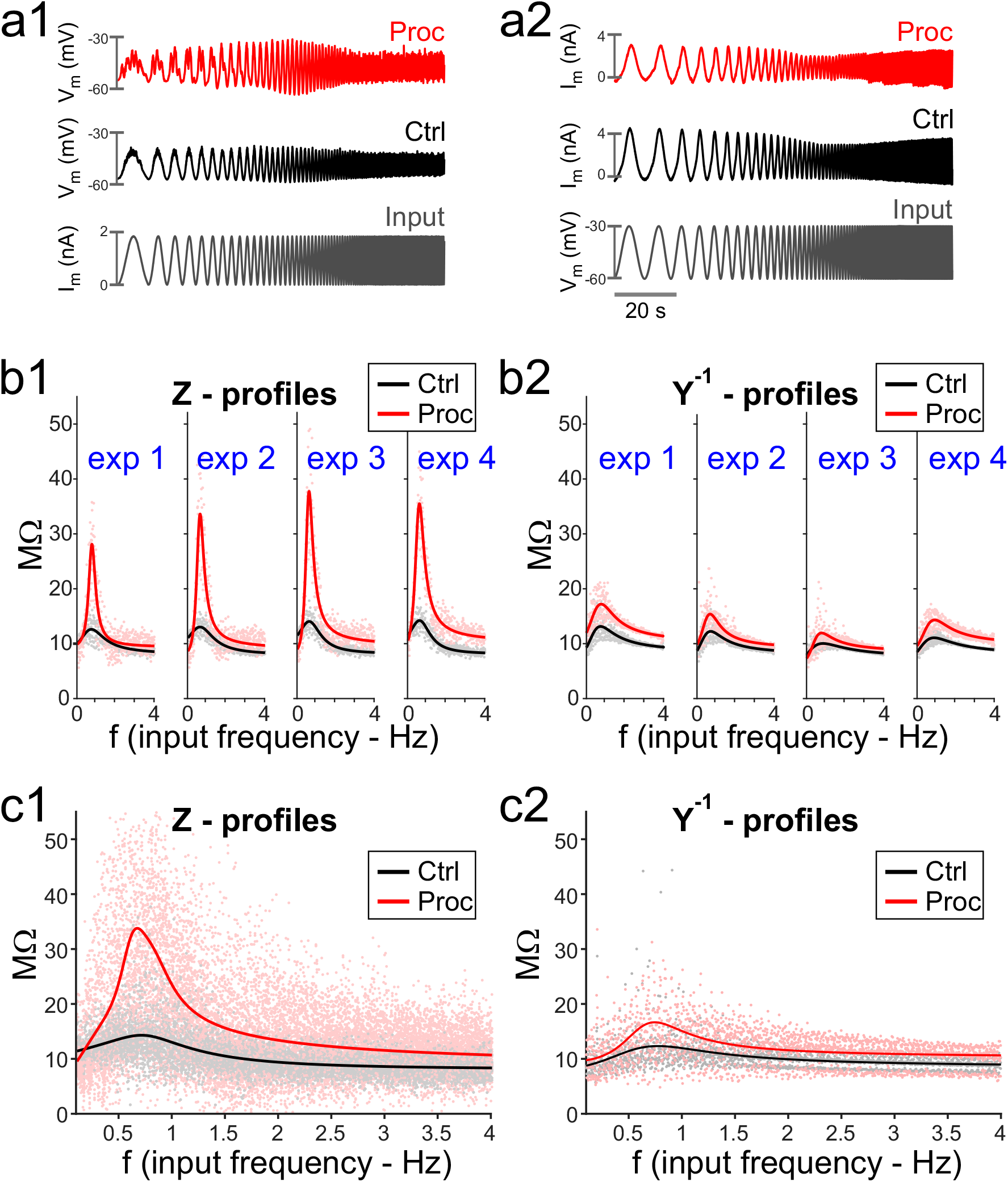
Voltage and current responses to chirp current and voltage inputs, respectively, for the biological PD neuron. **(a)** The membrane current response of the biological PD neuron, voltage clamped with a chirp function sweeping frequencies of 0.1 to 4 Hz and the voltage range of −60 to −30 mV. Responses are shown in control saline (Ctrl, **a1**) and in the presence of 1 *μ*M proctolin (Proc, **a2**). Proctolin activates the voltage-gated current *I_MI_*, which acts as an amplifying current. **(b)** The impedance [**b1**] and inverse admittance [**b2**] profiles, corresponding to the protocols shown in panel a, for individual experiments (1-4), in which the measurements were done in both I-clamp and V-clamp. Profiles shown are quadratic rational fits of each experiment. Dots show corresponding data point measurements for each experiment. (**c**) The average impedance [**c1**] and inverse admittance [**c2**] profiles corresponding to the experiments shown in **b**. Dots show all data point measurements.

## Discussion

Membrane potential (subthreshold) resonance has been studied in neurons using both the I-clamp and V-clamp techniques [5–54]. Studies that use the V-clamp technique to examine resonance, implicitly or explicitly assume the equivalence of the two methods (see, e.g., our own study [22]). Notably, despite the differences in dynamic complexity between both approaches, for linear systems, they produce equivalent results. However, subthreshold nonlinear effects, generated by the amplification processes resulting from nonlinear properties of ionic currents, may play significant roles in the communication of the neuronal subthreshold resonance properties to the spiking level [15, 16, 56–58]. These effects include not only the monotonically increasing dependence of the impedance profile with the input amplitude, but also the break of symmetry between the upper and lower envelopes of the voltage response that renders the impedance an imprecise predictor of the voltage response peak.

Our goal here is to point out and clarify some differences between nonlinear subthreshold responses to oscillatory inputs in I-clamp and V-clamp using a variety of neuronal models. It is important to note the primary dynamic difference between the two methods, which is the lower dimensionality of V-clamp as compared to I-clamp. This is the result of elimination, in V-clamp, of voltage as a dynamic variable. Because of this, complex responses to constant inputs such as nonlinear overshoots (depolarization or sags) and damped subthreshold oscillations that can be observed in I-clamp are not necessarily reflected in V-clamp. In contrast, the steady state responses of linear systems to constant inputs are completely equivalent in I-clamp and V-clamp, in the sense that they are the reciprocals of one another. Therefore, it is not surprising that for linear systems impedance and admittance are also reciprocals.

For quasilinear cases involving only resonant processes such as *I_h_* or *I_M_*, or their interplay with low levels of amplifying processes such as *I_Nap_* in which the system remains quasi-linear [57, 58], I-clamp and V-clamp still produce similar results for membrane potential resonance. However, the full equivalence between voltage and current responses in I-clamp and V-clamp breaks down for nonlinear systems. Even with simple nonlinearities, the differences between the two methods are relatively easy to see for constant inputs. For example, for the parabolic system (3)-(4), the steady state voltage response to constant *I* involves a square root with *I* inside the radical, while the steady state current response to constant values of *V* is a quadratic function of *V*. In order to understand the onset of the differences between I- and V-clamp we used asymptotic analysis (regular perturbation analysis) on weakly quadratic models. As a result, we concluded that effects of the nonlinearities are stronger on the *Z*-profiles than on the corresponding *Y*^−1^-profiles and that the effects of nonlinearities on the voltage equation are stronger on both profiles than if the same nonlinearities occur in the recovery variable equation, consistent with previous findings [55]. For time-dependent inputs, such as oscillations, these differences are somewhat more difficult to discern, since the equations are not analytically solvable. It is possible, however, to use methods of harmonic analysis to provide numerical approximations for the frequency response of nonlinear systems.

We used numerical methods to examine the differences between I- and V-clamp responses in additional nonlinear models. Our main findings are that the effects of nonlinearities are stronger when they are located in the voltage equation than in the recovery variable equations, and in the former case, the nonlinear amplifications are significantly stronger in I-clamp than V-clamp.

Finally, we examined the predictions of our analysis in an identified biological neuron, the PD neuron in the crab stomatogastric ganglion. The PD neuron shows resonance at a frequency of around 1 Hz, as measured both I- and V-clamp conditions [62–64]. Additionally, the neuropeptide proctolin activates a low-threshold voltage-gated inward current (*I_MI_*) in this neuron [66, 67], which has voltage-dependence and kinetics similar to the persistent sodium current *I_Nap_* and should therefore act as an amplifying factor for the resonance properties of this neuron. As predicted from our analysis, addition of proctolin only moderately increased the resonance peak of the inverse admittance profile measured in V-clamp, but the same treatment produced a large enhancement of resonance of the impedance profile in I-clamp. Biophysically, this difference could be explained by the fact that, in V-clamp, the regenerative properties of the amplifying current *I_MI_* are restrained by limitations on changes in the membrane potential.

The quantitative differences between I-clamp and V-clamp are present and similar in all nonlinear systems that we explored. Furthermore, the differences in the frequency-dependent responses measured in the two methods becomes greater as the structure of the nonlinear system, near the resting state, deviates from its linearization. However, in all cases, the existence of resonance was independent of whether the I-clamp or the V-clamp method was used. Because nonlinear systems could in principle produce a variety of responses depending on the details of the system and the amplitude of the input, it may be possible to imagine cases in which resonance only appears in I-clamp but not in V-clamp, or even vice versa. A full exploration of such a possibility requires simulations and analysis beyond the scope of our study and, we conjecture, would depend on the structure of the nonlinearities present in the system.

In conclusion, although V-clamp allows for better control of the experimental measurements when different conditions (modulators, synaptic effects, etc.) are compared (e.g. see [64]), in the presence of large nonlinear currents, such as regenerative inward currents that act as amplifying factors of resonance, measurements in I-clamp provide a more reliable characterization of the frequency-dependent responses of neurons.

## Acknowledgments

This work was partially supported by the National Science Foundation grants DMS-1313861, DMS-1608077 and National Institutes of Health grants MH060605 and NS083319.

## A Equivalence between the I-clamp impedance and the V-clamp admittance for linear systems

The I-clamp impedance and the V-clamp admittance are equivalent if the corresponding amplitudes are the reciprocal of one another and the corresponding phases have the same absolute value but different sign. Using the notation introduced in this paper, *Z*(*ω*) = *Y*^−1^(*ω*) and Φ(*ω*) = −Ψ(*ω*).

We illustrate this for the following linear system,

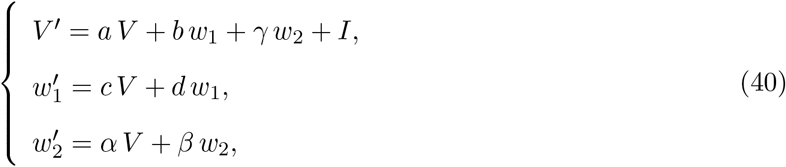

where the “prime” sign represents the derivative with respect to time *t* and *a, b, c, d, α, β* and *γ* are constants satisfying the condition that the eigenvalues of the characteristic polynomial for (40) with a constant value of *I* have non-positive real part. System (40) has the structure of the linearized conductance-based models [6, 68] for the voltage (*V*) and two gating variables (*w*_1_ and *w*_2_).

We assume

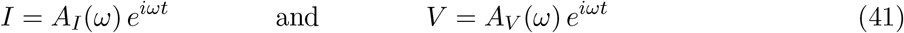

where *ω* is the frequency (a linear function of the input frequency *f*). In I-clamp *A_I_* = *A_in_* and *A_V_* = *A_out_*, while in V-clamp *A_I_* = *A_out_* and *A_V_* = *A_in_*. Typically, *A_in_* is independent of *ω*, but this need not be the case. Note that eqs. (40) are forced 3D and 2D linear systems in I- and V-clamp, respectively.

The I-clamp impedance and the V-clamp admittance are defined as

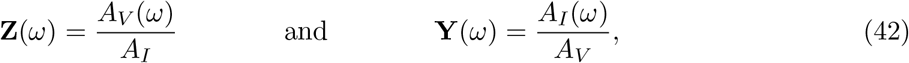

respectively, where **Z** and **Y** are complex quantities with amplitude (*Z* and *Y*, respectively) and phase (Φ and Ψ, respectively).

Alternatively, in I-clamp

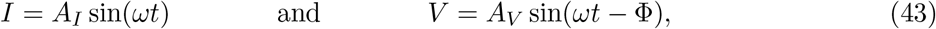

and in V-clamp

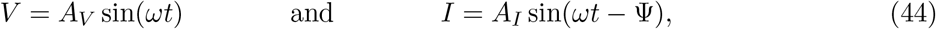

where *A_I_* and *A_V_* are real quantities. According to this formulation, *A_I_* = *A_in_* and *A_V_* = *Z*(*ω*) = |**Z**(*ω*)| in I-clamp, and *A_I_* = *Y*(*ω*) = |**Y**(*ω*)| and *A_V_* = *A_in_* in V-clamp.

The particular solutions (neglecting transients) of the second and third equations in (40) are given, respectively, by

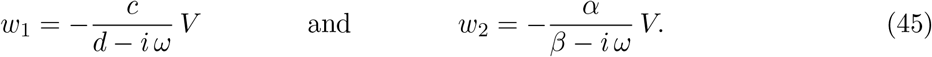

Substituting (45) into the first equation in (40) and rearranging terms yields

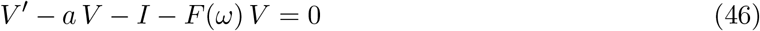

where

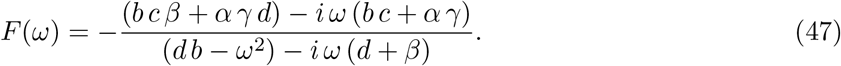

Substituting (41) into (46) gives the condition

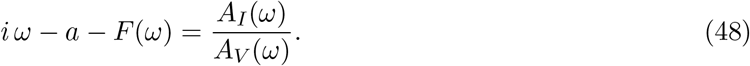

Therefore,

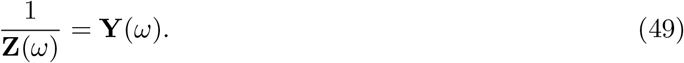

and

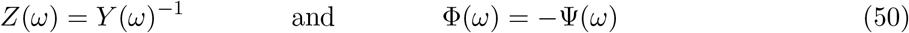

## B Solutions to oscillatory forced linear ODEs

### B.1 A system of two forced ODEs

Any system of ODEs of the form

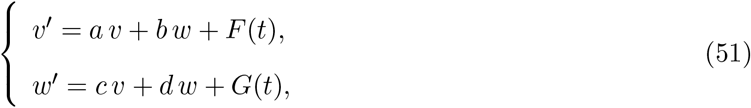

can be written as

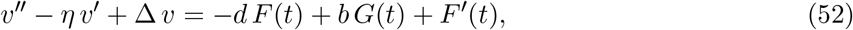

where

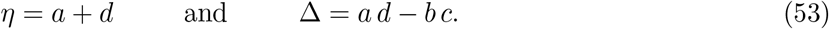

If *F*(*t*) and *G*(*t*) are linear combinations of sinusoidal and cosinusoidal function of the same frequency (*kω*), so there is the right-hand side of eq. (52). Therefore it suffices to solve

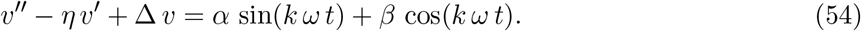

The solution of eq. (54) is given by

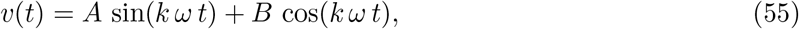

where

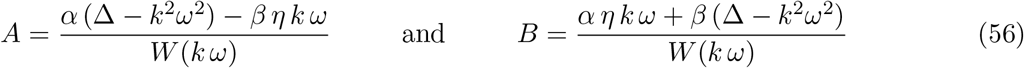

with

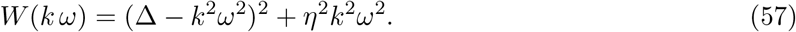

This solution satisfies

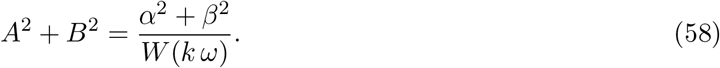

### B.2 A single forced ODE

The solution to any ODE of the form

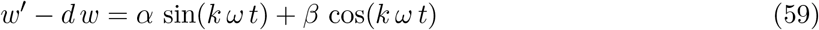

is given by

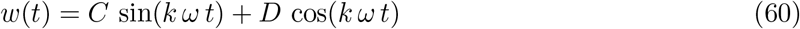

where

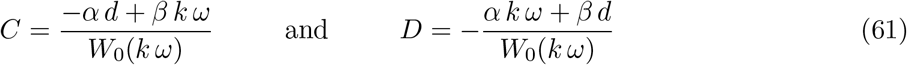

with

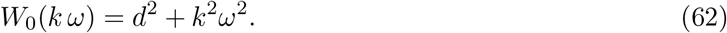

This solution satisfies

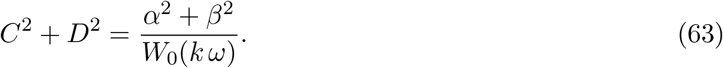

## C Linear systems receiving oscillatory inputs in I-clamp and V-clamp

### C.1 A linear system in I-clamp

System (51) with *F*(*t*) = *A_in_* sin(*ωt*) and *G*(*t*) = 0 can be written as

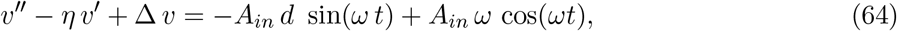

whose solution is given by (Appendix B.1)

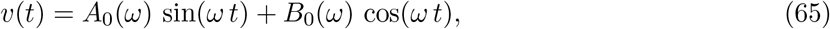

where

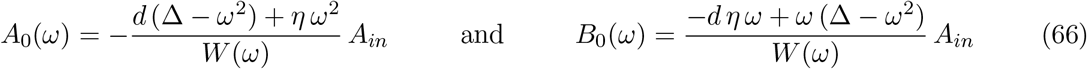

with *W*(*ω*) given by (57) with *k* = 1. From (58)

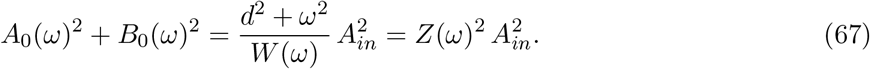

### C.2 A linear system in V-clamp

System (51) with *F*(*t*) = *I*, *v*(*t*) = *A_in_* sin(*ωt*) and *G*(*t*) = 0 can be written as

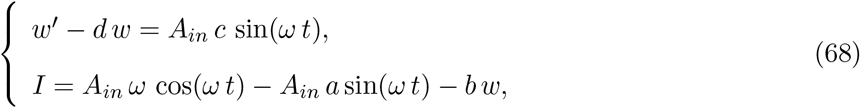

The solution to the first equation in (68) is given by (Appendix B.2)

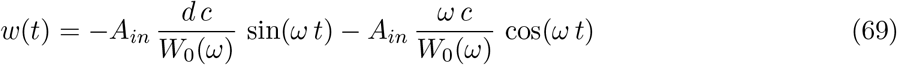

with *W*_0_(*ω*) given by (62) with *k* = 1. Substitution into the second equation in (68) yields

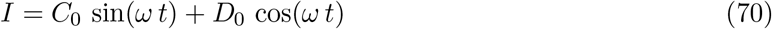

where

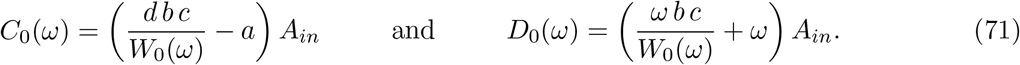

It can be shown that these constants satisfy

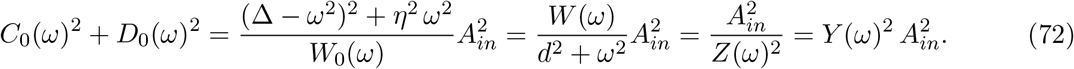

### C.3 A linearized conductance-based model in I-clamp

The solution to System (1)-(2) with *I*(*t*) = *A_in_* sin(*ωt*) (I-clamp) is given by (Appendix D)

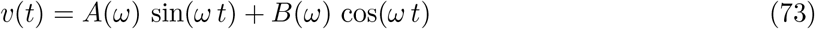

where

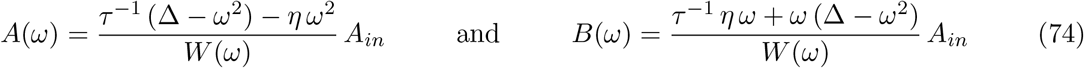

with

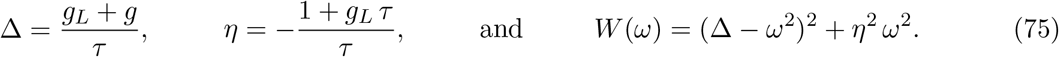

From (58),

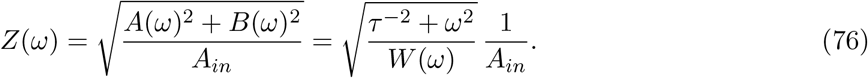

### C.4 A linearized conductance-based model in V-clamp

If, instead, we assume that *v*(*t*) = *A_in_* sin(*ωt*) (V-clamp), then the solution to eq. (2) is given by (Appendix D)

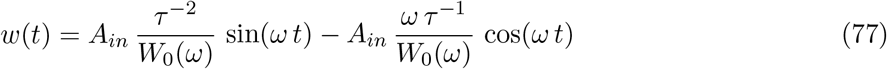

with

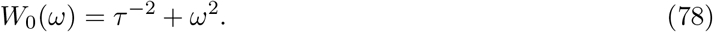

Substitution into the second equation in (68) yields

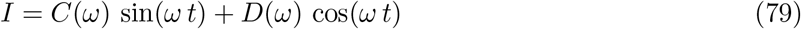

where

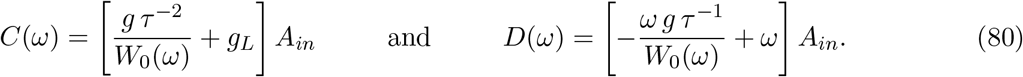

It can be easily shown that

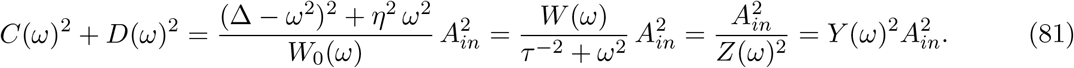

## D Weakly nonlinear forced systems of ODEs in I- and V-clamp: asymptotic approach

### D.1 Oscillatory input in I-clamp

We consider the following weakly perturbed system of ODEs

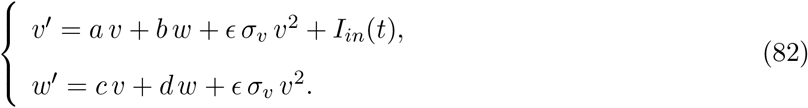

where

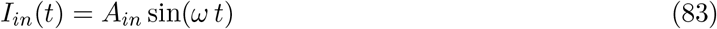

and *ϵ* is assumed to be small. We expand the solutions of (82) in series of *ϵ*

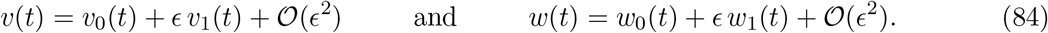

Substituting into (82) and collecting the terms with the same powers of *ϵ* we obtain the following systems for the 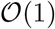 and 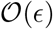 orders, respectively,

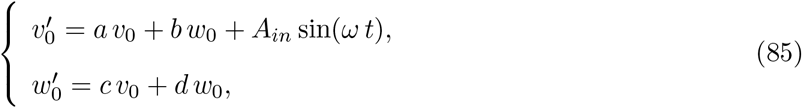

and

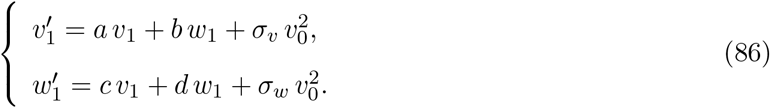

#### Solution to the 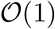 system

The solution to System (85) is given in Appendix C.1 with *v* substituted by *v*_0_.

#### Solution to the 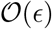 system

System (86) can be rewritten as

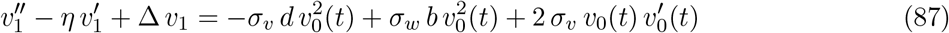

where

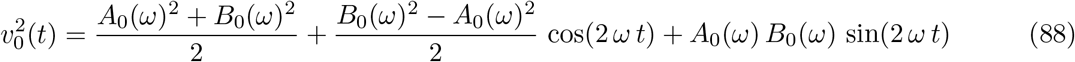

The solution to (87) is given (Appendix B.1) by

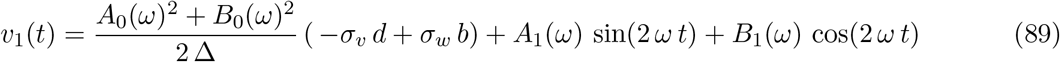

where

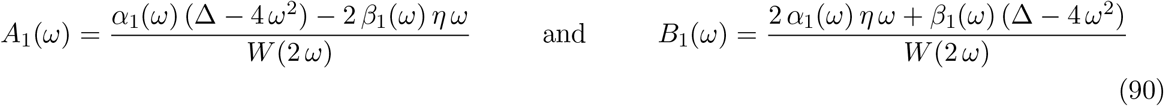

with *W*(2*ω*) given by (57) with *k* = 2,

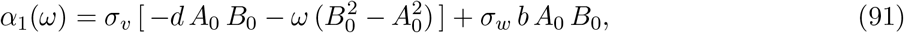

and

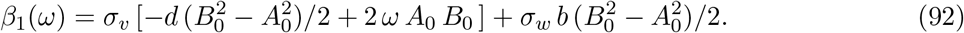

### D.2 Oscillatory input in V-clamp

We consider the following weakly perturbed system of ODEs

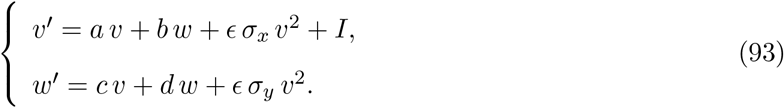

where

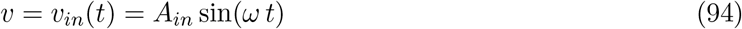

and *ϵ* is assumed to be small. We expand the solutions of (93) in series of *ϵ*

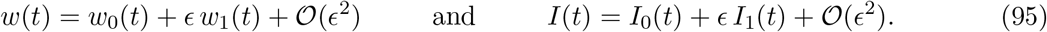

Substituting into (93) and collecting the terms with the same powers of *ϵ* we obtain the following systems for the 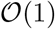 and 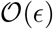 orders, respectively,

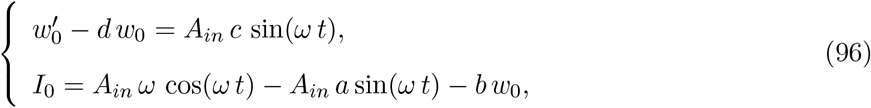

and

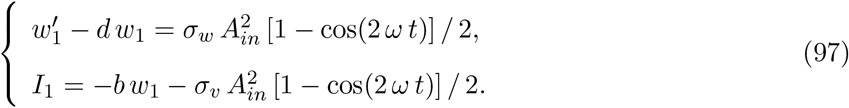

#### Solution to the 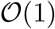 system

The solution to System (96) is given in Appendix C.2 with *w* and *I* and substituted by *w*_0_ and *I*_0_, respectively.

#### Solution to the 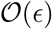 system

The solution to the first equation in (97) is given by (Appendix B.2)

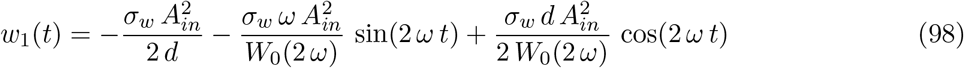

with *W*_0_(2*ω*) given by (62) with *k* = 2. Substitution into the second equation in (97) yields

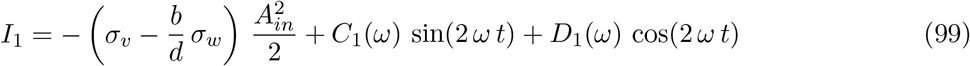

where

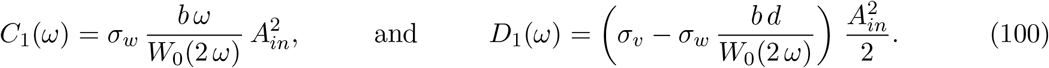

## E Asymptotic formulas for large values of *τ*

### E.1 Impedance zeroth-order approximation in I-clamp

For large enough values of *τ*, the coefficients of the solutions to the linear system (16) satisfy 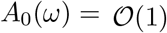 and 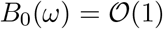, and

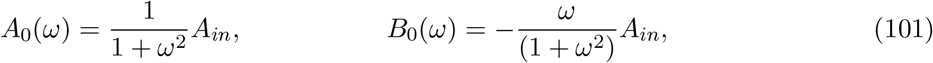

and

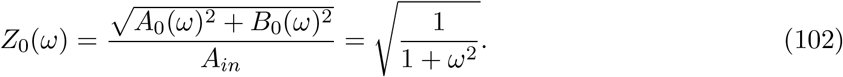

We begin with eqs. (74) and (75) and assume all other parameter values are 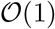. For large enough values of *τ* these quantities behave as follows

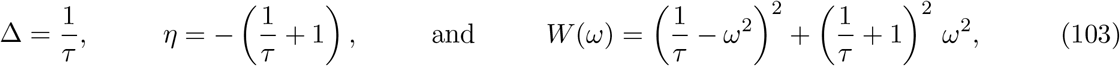

and

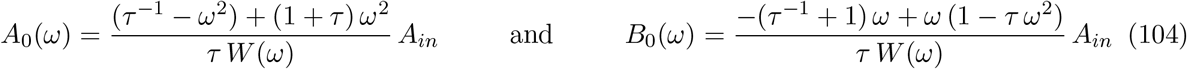

where

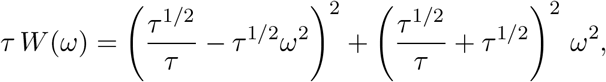

which can be reduced to

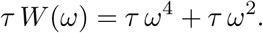

Substituting into (104) and rearranging terms yields (101) and 102.

### E.2 Admittance first-order approximation in V-clamp

For large enough values of *τ*

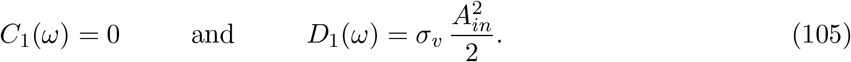

From (36) this implies that

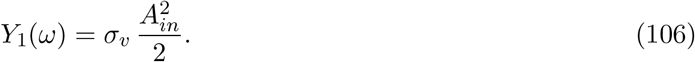

We begin with eqs. (37), for *C*_1_(*ω*) and *D*_1_(*ω*), and eq. (62) with *k* = 2 and *d* = −*τ*^−1^ for *W*_0_(2*ω*). Multiplication of the latter by *τ* and *τ*^2^ yields, respectively,

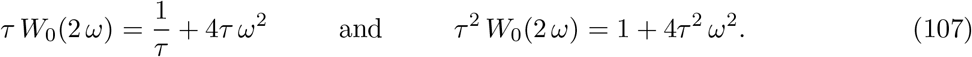

For large values of *τ*

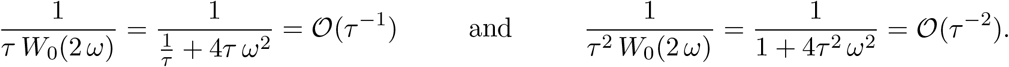

Therefore, for large enough values of *τ* in (37) we obtain (105).

### E.3 Impedance first-order approximation in I-clamp

For large enough values of *τ*

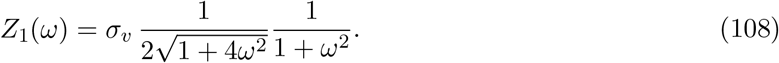

From (24) and (25) and the fact that 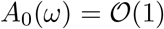 and 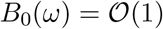 (Appendix E.1), it follows that for large enough values of *τ*

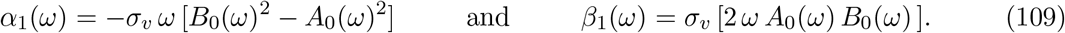

Substituting into (22) and rearranging terms we obtain

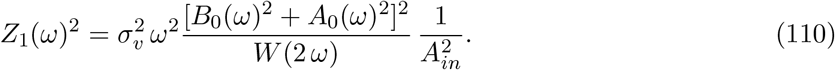

From (103) (and large enough values of *τ*)

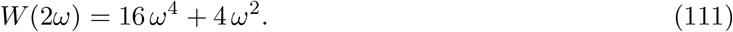

Substituting (111) and (101) into (110) we obtain (108).

